# Tirzepatide attenuates dopamine reward signaling and suppresses alcohol drinking and relapse-like behaviors in rodents

**DOI:** 10.1101/2025.08.26.672374

**Authors:** Christian E Edvardsson, Louise Adermark, Sam Gottlieb, Safana Alfreji, Thaynnam A Emous, Yomna Gouda, Annika Thorsell, Milica Vujičić, Cajsa Aranäs, Anna Benrick, Ingrid Wernstedt Asterholm, Marcelo F Lopez, Howard C Becker, Elisabet Jerlhag

## Abstract

Alcohol use disorder (AUD) remains a major public health problem, with few effective medications currently available. However, peptides of the gut-brain axis appear to offer promising therapeutic targets for AUD as they influence the mesolimbic reward circuitry. Here, we examined the effects of tirzepatide, a long-acting dual glucagon-like peptide-1 receptor (GLP-1R) and glucose-dependent insulinotropic polypeptide receptor (GIPR) agonist approved for diabetes and obesity, using behavioral assays, alcohol intake paradigms, and molecular analyses in rodents. First, tirzepatide effectively attenuated the rewarding properties of alcohol, measured through locomotor stimulation, conditioned place preference, and accumbal dopamine release. Subsequently, this GLP-1R/GIPR agonist dose-dependently reduced voluntary alcohol consumption, prevented binge and relapse-like drinking, and maintained efficacy during repeated administration. Finally, tirzepatide induced sustained synaptic depression in the lateral septum and further altered histone regulatory proteins in this region, suggesting a potential neural substrate for its effects. Moreover, the GLP-1R/GIPR agonist affected metabolic parameters including body weight, adipose tissue mass, hepatic triglycerides and circulating pro-inflammatory cytokines. Together, our findings suggest tirzepatide modulates alcohol-related behaviors through reward-related mechanisms while also affecting physiological consequences associated with long-term alcohol use. Given tirzepatide’s established clinical use and the consistency of effects observed here, these results support further investigation for treating AUD and associated complications.

**SIGNIFICANCE STATEMENT:** Existing treatments for alcohol use disorder show limited effectiveness, leaving patients without viable therapeutic options. We demonstrate that tirzepatide, a long-acting gut peptide-based drug already approved for diabetes and obesity, substantially reduces alcohol consumption and prevents relapse-like behavior across multiple preclinical models. Tirzepatide appears to work by influencing brain reward systems while simultaneously affecting metabolic complications common in alcohol disorders. Given tirzepatide’s clinical availability, these findings suggest repurposing a recently approved drug to tackle one of medicine’s more persistent treatment challenges.

## INTRODUCTION

Alcohol use disorder (AUD) remains a major public health challenge, contributing to substantial morbidity and mortality worldwide (1, 2). Despite available treatments, current medications show modest efficacy and are under-prescribed (3), highlighting the need for additional effective therapeutic approaches with alternative mechanisms of action. The neurobiological mechanisms underlying AUD involve the mesolimbic dopamine system, where alcohol-induced dopamine release in the nucleus accumbens (NAc) reinforces consummatory behaviors and contributes to the risk of developing AUD later in life (4-8). Long-term alcohol exposure leads to persistent neuroadaptations that disrupt mesolimbic system function, contributing to craving and relapse vulnerability (7-13). Research indicates that altered neuroplasticity and epigenetic mechanisms, including histone modifications, maintain these neuroadaptations (9-13). This complexity, combined with the limited success of existing therapies, suggests that current treatment approaches may be insufficient. Effective interventions might require strategies that address multiple interconnected systems influencing reward processing and addiction vulnerability.

In this context, gut–brain axis peptides have emerged as promising therapeutic candidates for AUD (14-16), given not only their apparent capacity to reduce alcohol intake (17) but also their wide-ranging physiological effects (18). The incretin hormones glucagon-like peptide-1 (GLP-1) and glucose-dependent insulinotropic polypeptide (GIP), traditionally recognized for their metabolic functions, also appear to influence central reward processing (14-16, 19, 20). Preclinical studies demonstrate that GLP-1 receptor (GLP-1R) agonists reduce alcohol consumption, likely by attenuating alcohol’s rewarding effects (21-28). Early clinical data from randomized trials and observational studies further demonstrates that GLP-1R agonists can reduce alcohol intake in humans (29-32). Building on these findings, clinical trials are now investigating these therapeutic applications more systematically, including studies examining incretin agonists for both alcohol consumption and alcohol-related disorders (ClinicalTrials.gov

identifiers: NCT06546384, NCT06409130, NCT07046819, NCT05891587, NCT05895643, NCT06015893, NCT05892432, NCT06939088, NCT06727331, and NCT06994338).

Recent advances have further introduced multi-receptor incretin agonists for diabetes and obesity treatment, with GLP-1R agonism serving as a central component (20). Given their apparent multiple modes of action, these compounds may address several aspects of AUD’s complex pathophysiology. Among these, tirzepatide, a long-acting dual GLP-1R/GIPR agonist approved for diabetes and obesity, shows enhanced therapeutic outcomes on cardiometabolic diseases compared to selective GLP-1R agonists (20, 33, 34). Tirzepatide’s clinical availability presents an opportunity to explore whether dual incretin agonists might offer advantages for AUD treatment. However, whether tirzepatide even affects alcohol consumption, and if so through what mechanisms, remains unexplored.

To address this knowledge gap, we conducted a systematic investigation of tirzepatide’s effects across multiple aspects of AUD using preclinical models. Our approach examined tirzepatide’s impact on alcohol-related reward processing, voluntary consumption, and binge and relapse-like drinking in both sexes. We also assessed tirzepatide’s influence on metabolic and inflammatory parameters, which are often dysregulated in AUD (35-37). To identify potential neural substrates underlying tirzepatide’s effects, we employed electrophysiological recordings across reward-related brain regions and conducted proteomic analysis of tissue samples from alcohol-consuming rats. Together, these studies allowed us to evaluate tirzepatide’s therapeutic potential while beginning to characterize what biological mechanisms might account for any observed effects.

## RESULTS

### Tirzepatide disrupts alcohol-induced behaviors and mesolimbic dopamine release in male mice

Alcohol consumption activates the mesolimbic dopamine system (6), producing locomotor stimulation, conditioned place preference (CPP), and dopamine release in the NAc. These behavioral and neurochemical changes underlie alcohol’s rewarding properties and contribute to the risk of developing AUD (5-8). Given tirzepatide’s potential to influence reward processing, we first tested whether acute tirzepatide administration (0.144 mg/kg, subcutaneously; SC) could alter these alcohol-induced (1.75 g/kg, intraperitoneally; IP) reward responses in male mice.

We first examined alcohol-induced locomotor stimulation. Tirzepatide alone had no effect on baseline locomotion compared to vehicle (P>0.999), while alcohol produced the expected locomotor activation (P<0.001) (F_3,32_=11.40, P<0.001; **Fig. 1A**). Tirzepatide pretreatment significantly blunted this alcohol-induced stimulation (P<0.001). In fact, locomotor activity in mice receiving both tirzepatide and alcohol was virtually indistinguishable from vehicle-only controls (P>0.999). We next examined tirzepatide’s influence on alcohol CPP. A control experiment verified that tirzepatide itself did not affect place conditioning when vehicle was paired with both compartments (t_14_=0.49, P=0.631; **Fig. 1B**). When alcohol was paired with one side, vehicle-treated mice developed clear preference for the alcohol-associated environment. Tirzepatide treatment markedly reduced this preference (t18=5.23, P<0.001; **Fig. 1B**). This guided us to examine another clinically relevant question. Recent evidence has shown that GLP-1R agonist exenatide reduces alcohol cue reactivity in NAc and septal regions of AUD patients (30), leading us to test whether tirzepatide might affect cue-induced place preference after prolonged abstinence. For this experiment we incorporated both environmental and olfactory cues in the CPP paradigm along with a 14-day abstinence period. While vehicle-treated mice retained strong preference for alcohol-associated contexts and cues on day 20, tirzepatide treatment substantially reduced this preference on the last test day (t_18_=4.13, P<0.001; **Fig. 1C**). These behavioral findings across locomotion, CPP, and cue-induced preference suggested that tirzepatide affects central reward processing mechanisms. We therefore employed microdialysis to investigate the neurochemical basis underlying these effects, examining alcohol-induced dopamine release in the NAc across two experimental paradigms. Systemic alcohol administration (**Fig. 1D**) produced pronounced elevation in NAc dopamine release compared to vehicle (treatment F_3,28_=57.47, P<0.001, interaction F_39,364_=17.42, P<0.001). Tirzepatide pretreatment substantially reduced this alcohol-induced dopamine response. Area under the curve analysis confirmed that tirzepatide alone did not affect baseline dopamine release (P>0.999), with no significant differences between vehicle controls and tirzepatide-alcohol treated mice (P=0.162). This effect moreover extended to alcohol-induced increases in dopamine metabolites 3,4-dihydroxyphenylacetic acid (DOPAC), 3-methoxytyramine (3-MT), and homovanillic acid (HVA), which tirzepatide similarly reduced (**Fig. S1A-C**). We also detected alterations in noradrenergic and serotonergic transmission (**Fig. S1D-G**), though these changes appeared less pronounced than the dopaminergic effects (**Fig. S1H**). To confirm these findings reflected tirzepatide’s influence on NAc dopamine responses specifically, rather than indirect systemic effects, we perfused alcohol locally through the probe (**Fig. 1E**). Local alcohol application evoked robust dopamine increases that systemic tirzepatide administration significantly attenuated (treatment F_1,14_=61.52, P<0.001, interaction F_13,182_=25.12, P<0.001), suggesting tirzepatide can modulate alcohol’s dopaminergic effects within the reward circuitry itself.

**Fig. 1.**
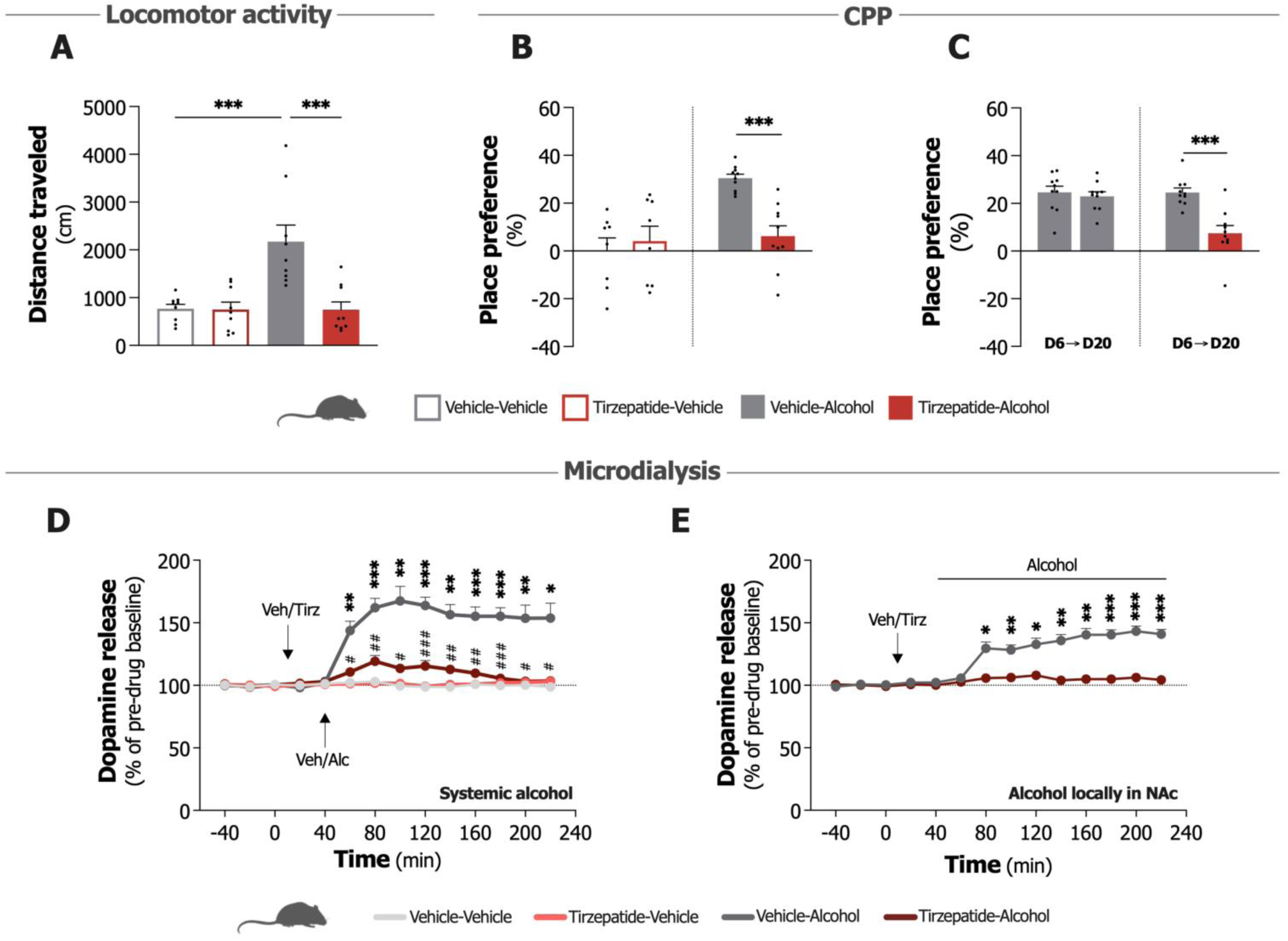
Impact of tirzepatide on alcohol-related reward behaviors and dopamine release in male mice. **A.** Tirzepatide (Tirz; 0.144 mg/kg) attenuates alcohol (Alc; 1.75 g/kg, IP)-induced locomotor stimulation (n=9/group, one-way ANOVA). **B.** Tirzepatide (0.144 mg/kg) treatment reduces the expression of alcohol (1.75 g/kg, IP)-induced conditioned place preference (CPP), without affecting CPP itself (n=8-10/group, unpaired t-test). **C.** On day 20 (D20) following a period of forced abstinence from day 6 (D6), tirzepatide (0.144 mg/kg) attenuates the expression of alcohol (1.75 g/kg, IP) and cue-induced CPP that persists in vehicle-treated mice (Veh), with the cue present on both testing days (n=10/group, unpaired t-test). **D.** Tirzepatide (0.144 mg/kg) significantly mitigates alcohol (1.75 g/kg, IP)-induced dopamine release in the nucleus accumbens (NAc) following systemic alcohol injection (n=8/group, repeated measures two-way ANOVA). **E.** Tirzepatide similarly blocks dopamine release when we perfused alcohol (300 mM) locally in the NAc (n=8/group, repeated measures two-way ANOVA). Data show mean ± SEM. *P<0.05, **P<0.01, ***P<0.001, ^#^P<0.05, ^##^P<0.01, ^###^P<0.001.

### Acute tirzepatide treatment dose-dependently reduces alcohol consumption in male and female rats

To further assess tirzepatide’s effectiveness across different alcohol drinking phenotypes and potential sex-specific effects, we conducted complementary experiments in male and female rodents. These experiments evaluated tirzepatide’s effects on voluntary alcohol consumption across multiple established paradigms: the intermittent access two-bottle choice model (38), the drinking in the dark (DID) model (39), and the alcohol-deprivation effect (ADE) model (40). We first assessed acute tirzepatide effects using two doses (0.048 and 0.144 mg/kg, SC) versus vehicle control on alcohol intake in the intermittent access paradigm to examine potential dose-dependent effects in both sexes. In male rats, the lower dose (0.048 mg/kg) showed minimal impact on alcohol, water, or food intake at 4 hours (**Fig. S2A-C**). By 24 hours, this dose significantly reduced alcohol intake (t_11_=7.81, P<0.001), food consumption (t_11_=6.07, P<0.001), and body weight (t_11_=9.60, P<0.001), while water intake remained unchanged (**Fig. S2D-G**). The higher dose (0.144 mg/kg) produced more pronounced effects, decreasing alcohol intake (t_11_=4.47, P=0.001) and food consumption (t_11_=4.75, P<0.001) while water intake showed an upward trend (t_11_=2.04, P=0.066) at the 4-hour timepoint (**Fig. S2H-J**). At 24 hours, this dose significantly reduced alcohol consumption (t_11_=6.14, P<0.001; **Fig. 2A**), decreased food intake (t_11_=18.90, P<0.001) and body weight (t_11_=15.30, P<0.001), while elevating water intake (t_11_=3.24, P=0.008; **Fig. S2K-M**). Direct comparison indicated that 0.144 mg/kg produced a significantly greater reduction in alcohol consumption (-51.7±6.3%) compared to the 0.048 mg/kg dose (-30.9±3.3%; t_22_=2.93, P=0.008), confirming dose-dependent effects (**Fig. S2N**).

**Fig. 2.**
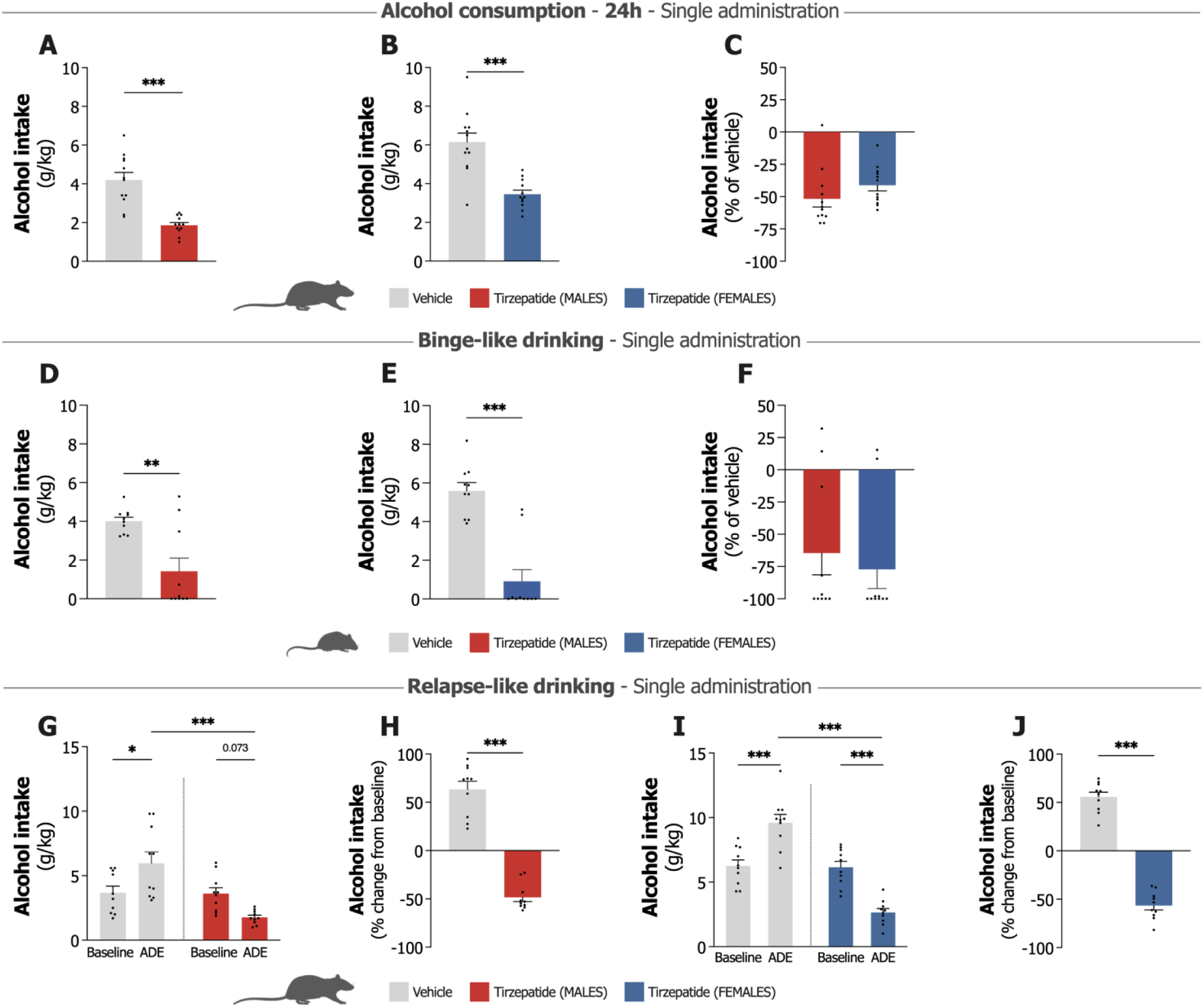
Effects of single administration of tirzepatide on alcohol intake, binge-like drinking and relapse-like drinking in male and female rodents. **A.** Single tirzepatide administration (0.144 mg/kg) significantly reduces 24-hour alcohol intake in male rats compared to vehicle (n=12, paired t-test). **B.** Female rats show similar reduction in 24-hour alcohol consumption following single tirzepatide treatment (0.144 mg/kg; n=12, paired t-test). **C.** Percent reduction in alcohol intake relative to vehicle demonstrates comparable tirzepatide efficacy (0.144 mg/kg) across both sexes with no statistically significant difference between males and females (n=12/group, unpaired t-test). **D.** Single tirzepatide administration (0.144 mg/kg) significantly attenuates binge-like drinking in male mice (n=10/group, unpaired t-test). **E.** Female mice also exhibit significant reduction in binge-like alcohol consumption following single tirzepatide treatment (0.144 mg/kg; n=10/group, unpaired t-test). **F.** Percent comparison against vehicle demonstrates comparable tirzepatide effectiveness (0.144 mg/kg) on binge-like drinking between sexes with no statistically significant difference between males and females (n=10/group, unpaired t-test). **G.** Vehicle-treated male rats exhibit significantly elevated alcohol intake during post-deprivation sessions compared to baseline consumption levels, whereas tirzepatide administration (0.144 mg/kg) effectively blocks this relapse-like drinking behavior (n=10/group, one-way ANOVA). **H.** Percent change from baseline in males demonstrates significant attenuation of relapse-like drinking behavior by tirzepatide (0.144 mg/kg; n=10/group, unpaired t-test). **I.** Vehicle-treated females demonstrate significantly elevated alcohol intake in post-deprivation sessions compared to baseline, whereas single-dose tirzepatide administration (0.144 mg/kg) inhibits this relapse-like alcohol drinking effect and further reduces alcohol intake compared to baseline levels (n=10/group, one-way ANOVA). **J.** Percent change from baseline in females shows tirzepatide efficacy (0.144 mg/kg) in preventing relapse-like drinking (n=10/group, unpaired t-test). Data show mean ± SEM. *P<0.05, **P<0.01, ***P<0.001.

Parallel studies in female rats revealed similar dose-dependent responses. The lower tirzepatide dose (0.048 mg/kg) produced minimal changes in alcohol, water, or food intake at 4 hours (**Fig. S3A-C**). At 24 hours, we found a trend toward reduced alcohol intake (t_11_=1.91, P=0.083), with significant decreases in food consumption (t_11_=3.78, P=0.003) and body weight (t_11_=2.92, P=0.014), while water intake remained unaffected (**Fig. S3D-G**). The higher dose (0.144 mg/kg) in females produced a trend toward lowered alcohol intake (t_11_=1.65, P=0.128), caused no change in food intake, and increased water intake (t_11_=2.71, P=0.020) at 4 hours (**Fig. S3H-J**). At 24 hours, this dose significantly reduced alcohol consumption (t11=6.96, P<0.001; **Fig. 2B**), diminished food intake (t_11_=10.24, P<0.001) and body weight (t_11_=7.12, P<0.001), while increasing water intake (t_11_=4.39, P=0.001; **Fig. S3K-M**). Direct dose comparison in females showed significantly greater alcohol intake suppression with the 0.144 mg/kg dose (-41.3±4.3%) compared to the 0.048 mg/kg dose (-11.7±5.7%; t₂₂=4.15, P<0.001), suggesting dose-dependent effects (**Fig. S3N**). We found no significant sex differences with the 0.144 mg/kg dose on alcohol intake between males (-51.7±6.3%) and females (-41.3±4.3%; t_22_=1.37, P=0.183; **Fig. 2C**). Alcohol consumption also returned to baseline levels 48 hours post-treatment in both sexes (**Fig. S4A-B**), indicating that the suppressive effect of an acute tirzepatide injection on alcohol intake did not extend beyond this timeframe.

### Tirzepatide attenuates binge-like drinking in male and female mice with dose-response effects similar to semaglutide

Given that binge-like alcohol consumption is a clinically relevant concern (41), we tested whether tirzepatide affects this drinking phenotype in male and female mice using the DID paradigm. Mice received either vehicle or tirzepatide (0.144 mg/kg, IP) one hour before dark onset, followed by four-hour alcohol access during the dark phase. Both male (t_18_=3.66, P=0.002; **Fig. 2D**) and female mice (t_18_=6.41, P<0.001; **Fig. 2E**) showed significant reductions in binge-like drinking, with no significant sex differences in efficacy: males (64.5±16.9%) versus females (77.2±14.9%) relative to vehicle (t_18_=0.56, P=0.581; **Fig. 2F**).

To further explore tirzepatide’s dose-response characteristics and compare them with semaglutide, given the latter’s emerging clinical evidence for alcohol reduction (29), we conducted additional studies using lower doses (0.001-0.072 mg/kg, IP). Both tirzepatide and semaglutide produced dose-dependent reductions in binge-like drinking across male (tirzepatide: F_6,63_=34.83, P<0.001; semaglutide: F_6,60_=21.36, P<0.001; **Fig. S5A-B**) and female mice (tirzepatide: F_6,62_=24.76, P<0.001; semaglutide: F_6,61_=50.45, P<0.001; **Fig. S5C-D**).

### Tirzepatide prevents relapse-like drinking in male and female rats

As preventing relapse remains one of the most persistent challenges in AUD treatment (9, 42, 43), we next tested whether tirzepatide could also affect relapse-like drinking behavior. Using the ADE model, which captures the temporary increase in alcohol consumption following forced abstinence (40), we assessed tirzepatide’s (0.144 mg/kg, SC) acute impact on this relapse-like response.

Following 10 days of alcohol deprivation, vehicle-treated male rats developed the expected ADE, with consumption rising significantly above baseline levels (F_3,36_=9.30, P<0.001; **Fig. 2G**). Tirzepatide treatment appeared to block this response entirely. Rather than showing increased drinking, treated rats exhibited a trend toward reduced alcohol intake compared to baseline (P=0.073). Direct comparison revealed substantial between-group differences: tirzepatide-treated males showed 48.3±4.4% reduction from baseline while vehicle-treated counterparts increased consumption by 63.5±8.3% (t_18_=11.90, P<0.001; **Fig. 2H**). Additionally, tirzepatide increased water intake while reducing both food consumption and body weight (**Fig. S6A-C**).

Female rats showed comparable responses. Vehicle-treated females developed robust ADE (F_3,36_=35.03, P<0.001; **Fig. 2I**), whereas tirzepatide treatment again prevented the rebound response and actually decreased alcohol consumption below baseline (P<0.001). Tirzepatide produced substantial prevention in females as well, with 56.5±4.6% reductions compared to 55.7±4.9% increases in vehicle controls (t18=16.80, P<0.001; **Fig. 2J**). As in males, tirzepatide also increased water intake while decreasing food consumption and body weight (**Fig. S6D-F**).

### Sustained effect of tirzepatide in reducing alcohol consumption with repeated administration in male and female rats

To address whether tirzepatide maintains effectiveness during repeated administration, an important consideration for any potential addiction therapy (44), we examined the effects of repeated tirzepatide (0.144 mg/kg, SC) or vehicle administration across six alcohol drinking days spanning two weeks following the baseline period.

Male rats receiving repeated tirzepatide showed sustained reductions in alcohol consumption (treatment F_1,18_=20.66, P<0.001, interaction F_5,90_=1.39, P=0.234; **Fig. 3A**), with consistent decreases compared to vehicle throughout the study period. Water intake remained unaffected by treatment (**Fig. S7A**), while tirzepatide significantly reduced both food consumption (treatment F_1,18_=78.47, P<0.001, interaction F_5,90_=12.67, P<0.001; **Fig. S7B**) and body weight (treatment F_1,18_=93.75, P<0.001, interaction F_5,90_=5.68, P<0.001; **Fig. S7C**) across the experimental timeline.

**Fig. 3.**
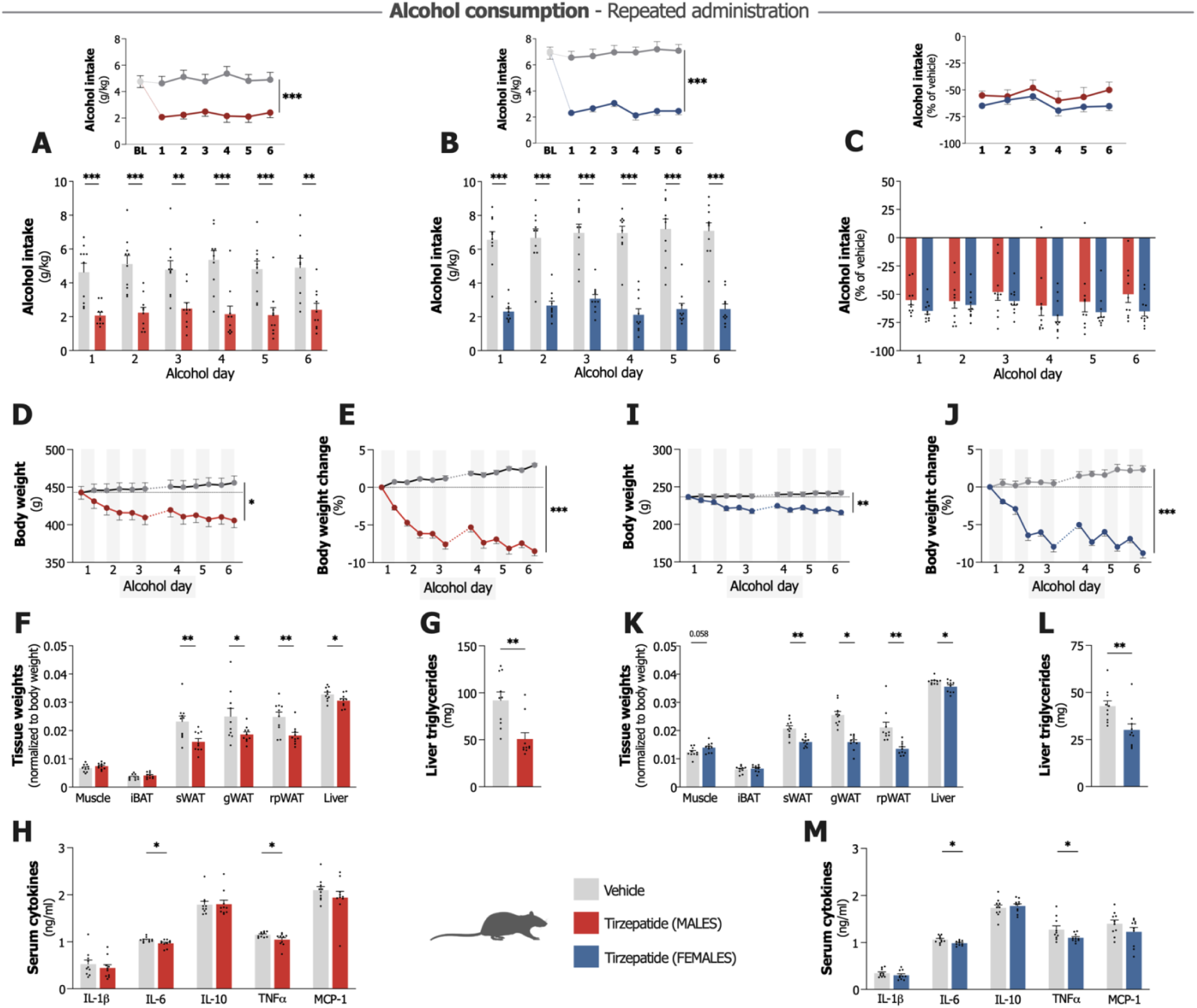
Effects of repeated tirzepatide administration on alcohol consumption, body composition and inflammation parameters in male and female rate. All experiments used n=10/group with repeated tirzepatide (0.144 mg/kg) or vehicle treatment. **A.** Tirzepatide significantly attenuates alcohol intake in male rats on all alcohol days compared to vehicle group, which shows similar baseline (BL) alcohol intake levels (repeated measures two- way ANOVA). **B.** Female rats show comparable reduced alcohol intake following tirzepatide administration, with consistent efficacy throughout all alcohol days (repeated measures two-way ANOVA). **C.** Alcohol intake relative to vehicle demonstrates comparable reducing effects across sexes with no statistically significant sex differences (repeated measures two-way ANOVA). **D-E.** Tirzepatide reduces body weight and percent body weight change in male rats compared to vehicle (repeated measures two-way ANOVA). **F.** Post-mortem tissue analysis in males shows significant reductions in subcutaneous inguinal (sWAT), gonadal (gWAT), and retroperitoneal (rpWAT) white adipose tissues and liver weight, while muscle (gastrocnemius) and intrascapular brown adipose tissue (iBAT) remain unaffected (unpaired t-tests). **G.** Tirzepatide significantly reduces hepatic triglyceride content in alcohol-drinking males (unpaired t-test). **H.** Treatment decreases interleukin (IL)-6 and tumor necrosis factor alpha (TNFα) serum levels, while IL-1 beta (β), IL-10 and monocyte chemoattractant protein-1 (MCP-1) remain unaffected (unpaired t-test). **I-J.** Female rats exhibit decreased body weight and percent body weight change following tirzepatide treatment (repeated measures two-way ANOVA). **K.** Tirzepatide significantly reduces white adipose tissue depots in females (unpaired t-tests). **L.** Tirzepatide significantly reduces hepatic triglyceride content in treated females (unpaired t-test). **M.** Treatment reduces IL-6 and TNFα serum levels in females, while levels of IL-1β, IL-10 and MCP-1 remain unaffected (unpaired t-test). Data show mean ± SEM with individual data points. *P<0.05, **P<0.01, ***P<0.001.

Female rats demonstrated similar sustained responses during repeated treatment. Tirzepatide produced comparable reductions in alcohol intake (treatment F_1,18_=84.59, P<0.001, interaction F_5,90_=1.24, P=0.296; **Fig. 3B**) that persisted throughout the two-week period. As observed in males, water consumption remained stable (**Fig. S7D**), while both food intake and body weight decreased significantly under tirzepatide administration (**Fig. S7E-F**).

Direct comparison revealed no significant sex differences in treatment response (sex F_1,18_=1.60, P=0.222, interaction F_5,90_=0.63, P=0.679), with females showing 63.4±4.1% reductions compared to 54.3±7.1% in males (**Fig. 3C**).

### Tirzepatide improves metabolic and inflammatory markers in alcohol-consuming rats across both sexes

Long-term alcohol use can lead to fatty liver disease, metabolic syndrome, and systemic inflammation, conditions that complicate treatment (35-37). GLP-1R agonists and tirzepatide have shown effects on metabolic liver disease and inflammatory processes (45-48). Given the notable body weight changes we observed during repeated treatment, we additionally explored whether tirzepatide might also affect these metabolic and inflammatory parameters in alcohol-consuming animals. Preliminary insights on these effects might inform whether tirzepatide could serve dual therapeutic roles, treating both alcohol use behaviors and the complications that often accompany AUD.

In male rats, body weight changes diverged significantly between treatment groups (treatment F_1,18_=6.39, P=0.021, interaction F_11,198_=95.70, P<0.001; **Fig. 3D**), with vehicle-treated rats gaining weight while tirzepatide-treated rats lost weight. Percentage body weight analysis showed more pronounced between-group differences (F_1,18_=181.80, P<0.001, interaction F_11,198_=97.35, P<0.001; **Fig. 3E**), with the tirzepatide group showing an average decrease of 8.5±0.6% and vehicle group an average increase of 3.0±0.2%. Tissue-specific analysis revealed selective effects of this weight reduction. Tirzepatide treatment did not significantly alter gastrocnemius muscle (t_18_=1.32, P=0.204; **Fig. 3F**) or intrascapular brown adipose tissue (iBAT) mass (t_18_=1.22, P=0.238) compared to vehicle. However, tirzepatide significantly reduced white adipose tissue (WAT) deposits across multiple locations: subcutaneous inguinal (sWAT; t_18_=3.10, P=0.006), gonadal (gWAT; t_18_=2.12, P=0.048), and retroperitoneal and perirenal (rpWAT; t_18_=1.619, P=0.005). Tirzepatide also decreased liver weight in treated males (t_18_=2.15, P=0.045; **Fig. 3F**), with accompanying reductions in hepatic triglyceride content (t_18_=3.70, P=0.002; **Fig. 3G**). Tirzepatide significantly reduced serum concentrations of pro-inflammatory cytokines interleukin-6 (IL-6; t_18_=2.49, P=0.023) and tumor necrosis factor alpha (TNFα; t_18_=2.17, P=0.043; **Fig. 3H**) in male rats, while levels of IL-1β, IL-10, and monocyte chemoattractant protein-1 (MCP- 1) remained unaffected.

Female rats exhibited comparable metabolic and inflammatory responses to tirzepatide treatment. Body weight changes showed similar divergence between treatment groups (F1,18=12.93, P=0.002, interaction F11,198=39.48, P<0.001; **Fig. 3I**), with tirzepatide inducing weight reduction while vehicle-treated females gained weight. Percentage body weight changes revealed an 8.8±0.6% decrease in tirzepatide-treated females compared to a 2.3±0.5% increase in vehicle-treated counterparts (F_1,18_=136.70, P<0.001, interaction F_11,198_=40.17, P<0.001; **Fig. 3J**). As observed in males, tirzepatide’s effects on body composition in females were tissue-specific. Tirzepatide preserved gastrocnemius muscle (t_18_=2.02, P=0.058; **Fig. 3K**) and iBAT mass (t_18_=0.59, P=0.564), while significantly reducing WAT deposits across all measured depots: sWAT (t_18_=4.49, P<0.001), gWAT (t_18_=6.38, P<0.001), and rpWAT (t_18_=3.84, P=0.001). Tirzepatide significantly reduced liver weight in treated females (t_18_=2.58, P=0.019; **Fig. 3K**), with concomitant decreases in hepatic triglyceride content (t_18_=2.99, P=0.008; **Fig. 3L**). The inflammatory profile mirrored that observed in males, with significant reductions in serum IL-6 (t_18_=2.62, P=0.017) and TNFα (t_18_=2.16, P=0.044; **Fig. 3M**) levels following tirzepatide treatment.

### Ex vivo electrophysiological recordings identified lateral septum as a possible target for tirzepatide’s neural effects

Before conducting a proteomic analysis of brain tissue from the repeated tirzepatide and long-term alcohol drinking experiment, where rats were sacrificed 24 hours after their final treatment, we performed electrophysiological screening to identify which reward-related circuits (9, 49) show detectable responses to tirzepatide treatment. Using alcohol-naïve male mice, we examined neural activity 24 hours after acute tirzepatide administration (0.144 mg/kg, SC) across several regions including NAc core and shell, medial prefrontal cortex (mPFC), dorsolateral and dorsomedial striatum (DLS/DMS), and lateral septum (LS).

Analysis of the LS recordings revealed that tirzepatide exposure 24 hours earlier produced a sustained suppression of evoked field potentials (F_1,56_=4.74, P=0.034; **Fig. 4A**), which was concomitant with a significant increase in paired-pulse ratio (t_40_=2.32, P=0.026; **Fig. 4B**), indicative of a decreased probability of neurotransmitter release. This suggests tirzepatide induced lasting presynaptic modifications within the LS region. In contrast, we observed no sustained differences when comparing tirzepatide-exposed mice with vehicle-treated mice in other reward-related circuits, including the mPFC (F_1,19_=0.01, P=0.924), DMS (F_1,34_=0.01, P=0.907), DLS (F_1,58_=1.06, P=0.308), NAc core (F_₁,₁₉_=0.26, P=0.617), or NAc shell (F_1,58_=0.48, P=0.490; **Fig. 4C-G**).

**Fig. 4.**
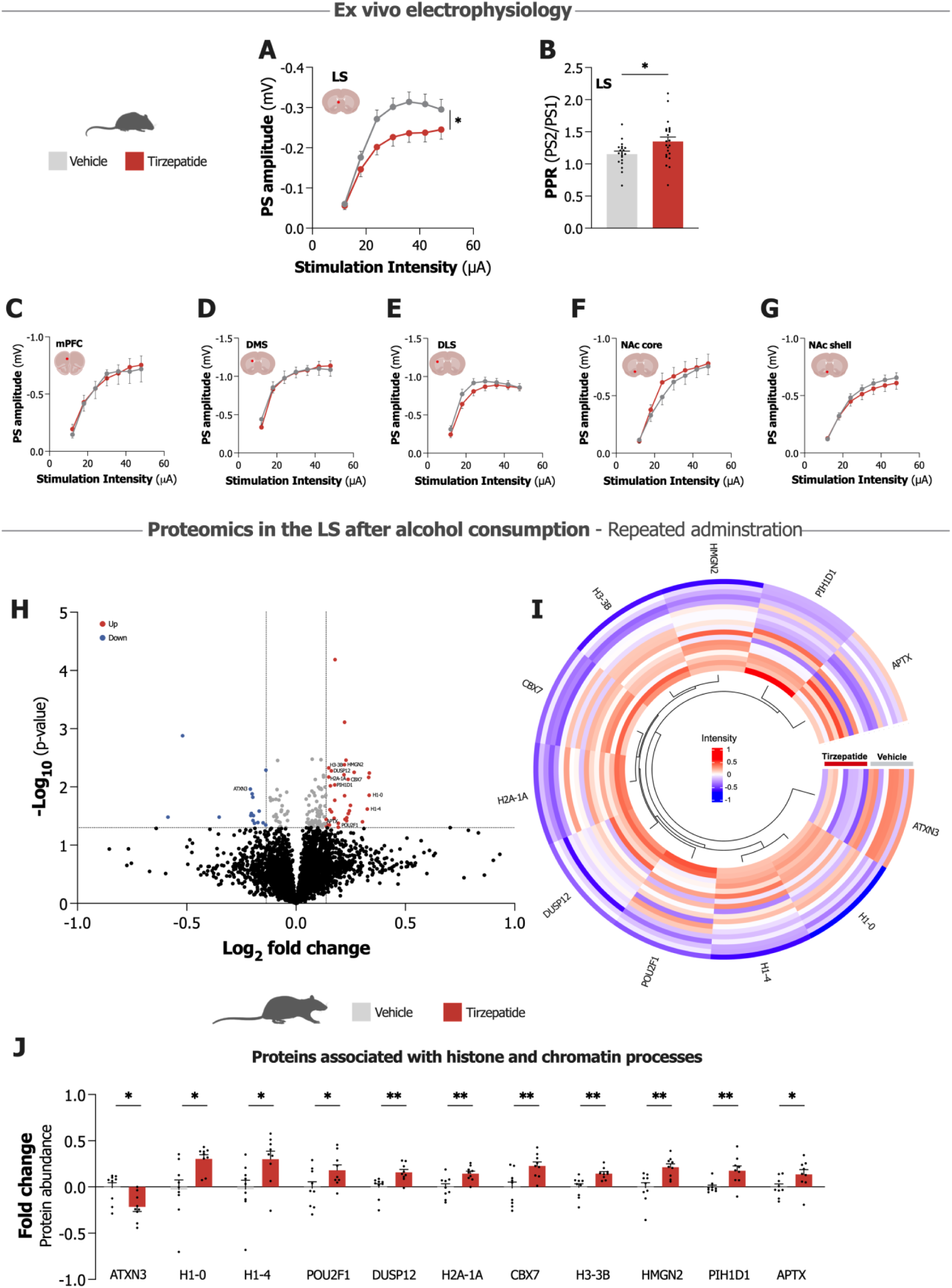
Electrophysiological effects of tirzepatide and proteomic alterations following tirzepatide administration in alcohol-exposed male rodents. **A.** Input-output curves derived from ex vivo brain slice field recordings from male mice demonstrate tirzepatide-induced reduction in population spike (PS) amplitude within the lateral septum (LS) (n=28-30/group, repeated measures two-way ANOVA). **B.** Paired-pulse ratio (PPR, PS2/PS1) measurements in the LS reveal significant effects in tirzepatide-treated mice compared to the vehicle group (n=20-22/group, unpaired t-test). **C-G.** Input-output curves in the medial prefrontal cortex (mPFC), dorsomedial striatum (DMS), dorsolateral striatum (DLS), nucleus accumbens (NAc) core, and NAc shell demonstrate region-specific electrophysiological responses, with no statistically significant differences observed in these regions following tirzepatide treatment (n=9-30/group, repeated measures two-way ANOVA). **H.** Volcano plot visualizes the proteomics data in the LS of alcohol-consuming male rats following repeated tirzepatide (0.144 mg/kg) administration, showing the differential protein expression profile. **I.** Circular heatmap visualizes differentially expressed proteins associated with histone and chromatin processes following repeated tirzepatide treatment in alcohol-drinking male rats. **J.** Quantification shows fold changes in specific proteins associated with histone and chromatin processes comparing tirzepatide treatment against vehicle, with significant differences observed in multiple proteins (n=9/group, Welch’s t-test, *p<0.05, **p<0.01).

### Proteomic analysis of the LS reveals chromatin regulatory proteins as potential targets of tirzepatide’s effects

Based on electrophysiology results identifying the LS as a region showing sustained responses to tirzepatide, we conducted a global quantitative proteomic analysis of the LS using tissue from male alcohol-drinking rats that received repeated tirzepatide treatment. We identified 4,359 distinct proteins using TMT mass spectrometry, with statistical analysis showing 51 proteins differentially expressed between tirzepatide and vehicle groups (**Fig. 4H**, **Table S1**). Tirzepatide upregulated 35 proteins and downregulated 16 proteins compared to vehicle.

Gene Ontology analysis identified several functional categories among the differentially expressed proteins (DEPs). We found proteins previously linked to alcohol consumption, including peroxisomal trans-2-enoyl-CoA reductase (PECR), midkine (MDK), acetyl-CoA acyltransferase 2 (ACAA2), ATP-binding cassette subfamily G member 2 (ABCG2), and reticulon 1 (RTN1). Tirzepatide also affected proteins involved in neurotransmission, such as solute carrier family 6 member 6 (SLC6A6), reticulon 3 (RTN3), proline-rich transmembrane protein 2 (PRRT2), alpha-aminoadipic semialdehyde synthase (AASS), and microtubule-associated protein 1A (MAP1A). We also detected changes in neuroinflammation-related proteins, including PECR, signal regulatory protein alpha (SIRPA), and leucine-rich repeat containing 14 (LRRC14).

The DEPs also included several histone proteins: H1-0, H1-4, H2A-1A, and H3-3B. Given that histone modifications and chromatin remodeling represent important epigenetic mechanisms involved in addiction pathophysiology (10, 11), we focused detailed analysis on GO terms related to histone and chromatin processes. Cross-referencing with the UniProtKB database (50) and existing literature revealed a total of 11 proteins associated with histone or chromatin regulatory functions, visualized as a heat map (51) in **Fig. 4I** (with references listed in **Table S1**). Statistical analysis showed significant differences between tirzepatide and vehicle groups in these 11 proteins: ataxin-3 (ATXN3; t_15.9_=2.88, P=0.011), histone H1-0 (t_10.7_=2.93, P=0.014), histone H1-4 (t_15.7_=2.50, P=0.024), POU domain class 2 transcription factor 1 (POU2F1; t_15.7_=2.14, P=0.049), dual specificity protein phosphatase 12 (DUSP12; t_15.5_=3.24, P=0.005), histone H2A-1A (t_14.2_=3.17, P=0.007), chromobox protein homolog 7 (CBX7; t_14.2_=3.12, P=0.007), histone H3-3B (t_12.5_=3.43, P=0.005), high mobility group nucleosome-binding domain-containing protein 2 (HMGN2; t_14.2_=3.41, P=0.004), PIH1 domain-containing protein 1 (PIH1D1; t_10.5_=3.19, P=0.009), and aprataxin (APTX; t_14.0_=2.22, P=0.043) (**Fig. 4J**).

## DISCUSSION

Here we demonstrate that tirzepatide, a dual GLP-1R/GIPR agonist, affects alcohol intake and alcohol-related responses across both sexes in rodents. Our findings reveal that tirzepatide attenuates alcohol-induced dopamine signaling, reduces alcohol consumption, and suppresses relapse-like behaviors. We also observed changes in metabolic and inflammatory parameters linked to alcohol use. Through electrophysiological and proteomic approaches, we identified the LS as what may be an important neuroanatomical substrate, with proteomic data suggesting potential epigenetic involvement in tirzepatide’s effects. These findings indicate that tirzepatide may represent a promising therapeutic candidate for AUD, potentially addressing both drinking behavior and the physiological consequences of alcohol intake that likely contribute to poor treatment outcomes and relapse vulnerability.

Our initial investigation examined tirzepatide’s effects on reward-related responses and revealed significant suppression of alcohol-induced dopamine processing. Tirzepatide consistently reduced alcohol-induced locomotor stimulation, place preference, and accumbal dopamine release in male mice across these paradigms. This modulation is noteworthy given that alcohol-induced dopamine release in the NAc appears to contribute to consummatory behaviors and likely represents a risk factor for AUD development (5-8). Perhaps most compelling was our observation that tirzepatide suppressed alcohol-induced dopamine release regardless of whether alcohol was administered systemically or perfused locally within the NAc itself. This suggests tirzepatide may directly influence the reward circuitry, though the precise mechanisms warrant further investigation. These findings align with previous work showing that GLP-1R agonists affect dopaminergic reward pathways across different substances of abuse (23-25, 27, 52-58).

The observed effects on reward processing appeared to translate into substantial reductions in alcohol-drinking behavior. Tirzepatide consistently decreased voluntary alcohol intake across both sexes in multiple experimental paradigms. Single administration produced dose-dependent reductions in alcohol consumption in rats while also suppressing binge-like drinking in mice, suggesting cross-species effectiveness. These observations extend what we and others have seen with GLP-1R agonists, where similar alcohol intake-suppressing effects have emerged across various preclinical models (21, 22, 25, 26, 28, 59-62).

Our comparison with semaglutide using similar dose ranges revealed similar acute effects on binge-like drinking for both compounds. This might suggest comparable therapeutic potential for both drugs when administered acutely, though it should be recognized that optimal dose ranges will likely differ between compounds in clinical settings, making direct dose comparisons challenging to interpret. A more intriguing observation was that the repeated tirzepatide administration appeared to maintain its suppressive effects more consistently than what we saw in our previous work with semaglutide (25), though we acknowledge this comparison spans different studies with inherent limitations. The clinical landscape looks increasingly promising. Early studies show that GLP-1R agonists reduce alcohol consumption in humans (29-32), suggesting our preclinical models might capture clinically relevant mechanisms. Cross-species translation always demands careful interpretation, though. Even more encouraging, a recent case-control study found reduced alcohol consumption in obese patients receiving tirzepatide (63), providing additional support for potential therapeutic applications. While these observations emerge from metabolic treatment contexts rather than addiction-focused studies, they provide what appears to be real-world validation of our experimental findings. Since tirzepatide and other GLP-1R agonists already have clinical approval for type 2 diabetes and obesity, a solid foundation exists for further AUD investigation. This practical advantage appears to have facilitated current clinical initiatives, with multiple studies now exploring incretin agonists potential as addiction treatment.

Our findings on relapse-like behaviors add further relevance to these clinical applications. Relapse remains a clinical challenge in AUD and effective treatment options remain scarce (9, 42, 43). We found that tirzepatide blocks ADE, a model of relapse-like drinking (40), which aligns with our earlier work with liraglutide and semaglutide (25, 59). Tirzepatide also attenuated cue-induced place preference following forced abstinence, suggesting possible effects on drug memory and cue reactivity processes (42, 43). Existing literature indicates that GLP-1R agonists also suppress reinstatement for cocaine (56), nicotine (57), and opioids (58), further supporting a potential effect on relapse-like behaviors. Furthermore, our findings seem translationally promising, given recent reports that semaglutide reduces alcohol cravings in humans (29) and that exenatide diminishes cue reactivity in AUD patients (30). Both anecdotal and research-based observations also support this interpretation, with reports of reduced cravings for food and substances during GLP-1R agonist treatment (63-65). As both cravings and environmental cues often precipitate relapse (42, 43), our findings further support the emerging evidence that incretin agonists could help mitigate relapse risk.

The repeated administration study also revealed effects on metabolic and inflammatory parameters that warrant further investigation. Given planned clinical trials exploring tirzepatide and semaglutide for alcohol liver disease and AUD patients with metabolic comorbidities (ClinicalTrials.gov identifiers: NCT06546384, NCT06409130, and NCT07046819), these findings could take on added significance. While incretin agonists demonstrate established benefits in metabolic and inflammatory conditions (45-48), their effects in long-term alcohol-consuming populations remain less well characterized. The question is whether they maintain these effects when alcohol consumption is involved. Our findings suggest they might. Repeated tirzepatide treatment reduced body weight, adipose tissue mass, liver weight, and hepatic triglyceride content, and also decreased pro-inflammatory cytokines (IL-6, TNFα) in both male and female alcohol-consuming rats. These observations suggest tirzepatide may offer therapeutic potential for alcohol-related complications such as fatty liver disease, a finding with clear implications for patients managing concurrent AUD and metabolic disorders. However, clinical studies specifically in alcohol-using populations will be necessary to validate these applications.

Moreover, our alcohol consumption studies demonstrated generally consistent effects across sexes, though future work should examine whether subtle sex-specific mechanisms exist at molecular and circuit levels.

Given the consistent effects across sexes, our subsequent investigations focused on male rodents to identify potential brain regions mediating tirzepatide’s effects on alcohol-related responses. We started with electrophysiological recordings across several reward-related circuits. This approach led us to the LS, where we observed the most pronounced changes in neural activity, including increased paired-pulse ratios suggesting modified presynaptic neurotransmitter release probability. These LS findings seemed particularly significant given the region’s established role in reward processing and drug-seeking behaviors (49, 66, 67). Several factors made this discovery especially compelling. The LS sits strategically near the subfornical organ, a circumventricular structure that might serve as an access point for peripherally administered incretin agonists (68). It also maintains direct connections to the NAc and ventral tegmental area (49, 66, 67), positioning it to potentially influence mesolimbic dopamine signaling (69). Both GLP-1R (70, 71) and GIPR (72, 73) are expressed in the LS, and peripherally administered GLP-1R agonists can reach this region (21, 74, 75). A recent clinical study have further shown that these drugs can reduce septal cue reactivity to alcohol in AUD patients (30). Previous preclinical work has also demonstrated LS involvement in GLP-1R-mediated reward processing for both alcohol (21, 24) and cocaine-related behaviors (52, 76), as well as regulation of alcohol-induced dopamine release in the NAc (24). This convergent evidence led us to hypothesize that the LS might serve as a neuroanatomical substrate mediating tirzepatide’s effects on mesolimbic dopamine signaling and alcohol-related responses.

To further explore potential LS-related molecular mechanisms, we conducted a proteomic analysis of LS tissue from alcohol-consuming male rats from the repeated tirzepatide treatment experiment. This revealed differential expression of several histone and chromatin regulatory proteins, including specific histone proteins (H1-0, H1-4, H2A-1A, H3-3B). These findings seemed noteworthy because histones regulate gene expression and serve as essential components of chromatin architecture (10, 13). Given that alcohol exposure can cause histone modifications, such as acetylations and methylations, which may contribute to addiction pathophysiology (10, 11, 77, 78), our results suggest tirzepatide might influence epigenetic mechanisms involved in alcohol drinking behaviors. Recent clinical research with semaglutide in obesity has identified similar proteomic changes in blood samples, including proteins linked to substance use disorders, such as histones (79). These findings hint that GLP-1R-based therapeutics may engage epigenetic mechanisms, tentatively explaining their broad efficacy across multiple disorders. However, future studies will need to investigate these potential mechanisms more thoroughly.

Our experimental approach provides solid evidence for tirzepatide’s therapeutic potential, though several important considerations deserve attention. We were careful to exclude potential confounding factors like sedation, anhedonia, and malaise since tirzepatide did not affect baseline locomotor activity, dopamine levels per se, or kaolin consumption. Even so, we did not specifically assess anxiety-like behaviors or taste aversion, which might influence alcohol-related responses. Our proteomic analysis focused on just one brain region, limiting our ability to understand broader molecular changes across reward circuits. An important mechanistic question also remains unanswered: did tirzepatide directly cause the molecular changes we observed, or did they simply result from reduced alcohol intake? This distinction seems crucial for understanding how tirzepatide actually works long-term. Targeted experimental designs could help address this to better understand the molecular effects we are seeing in the LS. Future studies should also explore the brain-circuit connectivity between LS GLP-1R/GIPR pathways and other regions since this could clarify which specific neural circuits contribute to tirzepatide’s effects on reward processing and alcohol consumption. It would also prove valuable to directly compare tirzepatide versus semaglutide within the same study to understand their different mechanisms. Another limitation worth noting: several experiments used only males, which probably limits our insights into sex-specific mechanisms. Even with these considerations, our findings suggest that tirzepatide influences the reward circuitry and may serve as a therapeutic candidate for AUD and related alcohol-related conditions.

In summary, our findings indicate that tirzepatide influences alcohol-related responses in ways that appear to have clinical potential. Tirzepatide consistently reduced alcohol intake across different drinking paradigms and both sexes without signs of tolerance development. Perhaps more significantly, tirzepatide’s effects on relapse behaviors suggest it might help decrease relapse vulnerability, a finding that could prove important for therapeutic applications. The mesolimbic reward effects offer insights into tirzepatide’s possible mode of action. Our data suggest tirzepatide influences dopaminergic reward processes to suppress alcohol-drinking behaviors. The LS findings add another piece to this puzzle, offering some initial clues about a neural substrate where dual incretin receptor signaling might exert these effects. These results, combined with tirzepatide’s existing clinical approval, position this dual incretin agonist as a promising therapeutic candidate for AUD and alcohol-related diseases, that warrants clinical investigation.

## METHODS AND MATERIALS

### Animals

Adult male NMRI mice (8-10 weeks old, 25-30 g; Charles River, Sulzfeld, Germany) were used for locomotor activity, CPP, microdialysis, food intake, and electrophysiological studies. Binge-like drinking paradigms employed adult male and female C57BL/6J mice (8-10 weeks old, 25-30 g; Jackson Laboratories, Bar Harbor, ME, USA). Intermittent access two-bottle choice alcohol studies utilized adult male and female Rcc/Han Wistar rats (8-9 weeks old, 180-250 g; Envigo, Horst, Netherlands), with tissue collected post-mortem for molecular analyses. The three rodent strains were selected based on their established responsivity to alcohol and gut-brain peptides (22-24, 80). Animals were group-housed upon arrival and acclimated for at least one week under standardized conditions (12/12-hour light/dark cycle, 20°C, 50% humidity) with ad libitum access to standard chow (Harland Teklad Rodent Diet #2916 & 2918, Madison, WI, USA) and water. Animals used for microdialysis and alcohol intake studies were subsequently single-housed after surgery or at the start of the alcohol baseline period to prevent implant damage and allow for individual consumption measurements. Behavioral and microdialysis experiments were conducted during the light phase when stimulation effects are more pronounced, with 60-minute habituation to the testing environment. Alcohol intake studies were performed during dark and light phases for rats and exclusively during the dark phase for mice, when drinking behavior is heightened. All experiments received approval from the Ethics Committee for Animal Research in Gothenburg, Sweden (ethical permits: 4685/23, 3348/20, 3276/20) or the Institutional Animal Care and Use Committee at the Medical University of South Carolina, USA. Studies adhered to the NIH Guide for the Care and Use of Laboratory Animals, ARRIVE guidelines, and 3Rs principle.

### Drugs

For behavioral and neurochemical experiments, alcohol (95% Ethanol, Solveco; Stockholm, Sweden or Warner Graham Co., Cockeysville, MD, USA) was diluted with vehicle (0.9% NaCl) to a 15% (w/v) solution and administered IP at 1.75 g/kg, 5 minutes prior to testing. Microdialysis experiments required local alcohol administration, achieved by diluting alcohol in modified Ringer’s solution (140 mM NaCl, 1.2 mM CaCl₂, 3.0 mM KCl, and 1.0 mM MgCl₂, Sigma-Aldrich, Darmstadt, Germany) to 300 mM, corresponding to approximately 50-60 mM outside the probe in the NAc (81). These alcohol doses and concentrations were selected for their established ability to stimulate the mesolimbic dopamine system with reproducible effects (22-24). Alcohol drinking studies employed a 20% (v/v) solution prepared with tap water.

Tirzepatide (LY3298176 HCl, MedChemExpress, Sollentuna, Sweden) was dissolved in vehicle (40 mM Tris-HCl, pH 8.0) and administered SC 30 minutes before behavioral testing or alcohol exposure. Dose selection was guided by initial dose-response studies (0.048, 0.144, 0.240, and 0.336 mg/kg) that evaluated effects on locomotor activity and food/kaolin intake. While no dose altered baseline two-hour locomotor activity or gross behavior (**Fig. S8A-D**), dose-dependent decreases in food intake and body weight appeared at 24 hours without affecting kaolin or water intake (**Fig. S9A-D**). Based on these findings, 0.144 mg/kg was selected for subsequent experiments to achieve consistent effects 30 minutes post-administration. The lower dose (0.048 mg/kg) was additionally tested in acute alcohol drinking studies to evaluate dose-dependent effects. Binge-like alcohol drinking experiments compared lower tirzepatide doses (0.001, 0.003, 0.009, 0.018, 0.036, and 0.072 mg/kg) with corresponding doses of semaglutide (MedChemExpress). Both compounds were administered IP one hour before dark onset in these studies. To validate this methodological variation, we compared SC versus IP administration effects on food intake for both tirzepatide (**Fig. S9A-D**) and semaglutide at a dose previously shown to attenuate alcohol-related behaviors (25) (**Fig. S9E-H**). Both administration routes produced comparable effects (**Fig. S9A-H**).

### Locomotor activity

Horizontal and vertical activity were recorded in six sound-attenuated, ventilated, and dimly lit (3 lx) locomotor boxes (42×42×20 cm; Open Field Activity System, Med Associates Inc., Georgia, VT, USA). Movement was detected via a two-layered infrared photobeam grid and recorded using Activity Monitor software (Version 7, Med Associates Inc.) (22-24). Male mice underwent 60- minute habituation in the test arena before first receiving tirzepatide treatment and then 30 minutes later an alcohol injection. Activity recording began 5 minutes after the final injection and continued for 60 minutes.

### Conditioned place preference

CPP experiments employed four two-chambered arenas (50×24×24 cm, custom-made, University of Gothenburg, Gothenburg, Sweden) under dim lighting (3 lx), with chambers distinguished by distinct tactile and visual cues (22-24). The protocol was conducted in male mice, beginning with a 20-minute pre-test (day 1) to assess initial place preference following vehicle injection. Conditioning sessions (days 2-5, 20 minutes each) followed a biased design, pairing alcohol with the least preferred chamber and vehicle with the preferred chamber. Daily sessions included one alcohol injection and one vehicle injection in a balanced design, alternating between morning and afternoon. On test day (day 6), mice received tirzepatide or vehicle before place preference monitoring for 20 minutes. A control experiment was conducted to assess tirzepatide’s effect on CPP independently of alcohol, following identical procedures but employing vehicle injections in both chambers during conditioning. A third experiment investigated tirzepatide’s influence on cue-induced place preference following forced abstinence, using the same biased alcohol paradigm with added neutral-valence olfactory cues (82, 83). Caraway odor (S-carvone, Sigma-Aldrich) was paired with the alcohol chamber and mineral oil (Sigma-Aldrich) with the vehicle chamber. Odorants (one drop) were applied to filter paper in perforated plastic tubes positioned at chamber tops. This protocol included a first test day (day 6) where mice received vehicle before a 20-minute test with olfactory cues present. Following two weeks of forced home cage abstinence, mice were divided into equal groups based on first test day results and received tirzepatide or vehicle before a second 20-minute test (day 20) with olfactory cues present. All experiments were analyzed using Observer XT software (Version 15, Noldus, Wagenegen, Netherlands). CPP expression was calculated as the difference in percentage of total time spent in the drug-paired compartment between pre-test and test sessions.

### Microdialysis

An I-shaped microdialysis probe (20 kDa cut-off membrane with 1 mm exposed length, HOSPAL, Gambro, Sweden) was surgically implanted in the NAc shell four days before experiments as previously described (22-24). Mice were anesthetized with isoflurane (Baxter, Apoteket AB, Gothenburg, Sweden), placed in a stereotaxic frame, and maintained on a heating pad. Local anesthesia (Xylocaine with adrenaline, 10 mg/ml, 5 μg/ml; Pfizer Inc, Apoteket AB, Gothenburg, Sweden) was applied at the incision site. Carprofen (Rimadyl®, 5 mg/kg, AstraZeneca, Apoteket AB, Gothenburg, Sweden), 0.9% NaCl, and Viscotears were administered for pain management, rehydration, and eye protection. After exposing the skull, holes were drilled for the probe and anchoring screws. The probe was secured with dental cement (DENTALON® Plus, Agntho’s AB, Lidingö, Sweden). On experiment days, the probe was connected to a pump and perfused with Ringer’s solution at 1.6 μl/min. After a two-hour equilibration period, samples were collected at 20-minute intervals throughout the experiment. Following baseline measurements (minutes -40 to 0), tirzepatide or vehicle was administered at minute 10. Thirty minutes later (minute 40), alcohol was either injected systemically (Experiment 1) or perfused through the probe for the remainder of the experiment (40-220 minutes, Experiment 2). Nine additional samples were collected following alcohol exposure. Probe placement was verified histologically using a brain atlas (84), and only data from correctly placed probes without hemorrhage were included in analyses (**Fig. S10A-B**). Microdialysate samples were analyzed using HPLC with electrochemical detection, as described before (24). Changes in dopamine and other monoamines and their metabolites were calculated as percentages of the mean of three baseline values before tirzepatide/vehicle treatment. The area under the curve following alcohol exposure (40-220 minutes) was additionally calculated for further analysis.

### Intermittent access two-bottle choice and the alcohol deprivation effect

This paradigm provided rats with alcohol and water access during three 24-hour sessions weekly (Monday, Wednesday, Friday), with water-only access on intervening days (24, 38). Bottles were switched at dark-phase onset, while food and water remained continuously available. Rats underwent an 8-week baseline period before experimental interventions, during which alcohol, water, and food intake were measured daily and body weight recorded weekly. Following the baseline period, rats were allocated to treatment groups with matched baseline alcohol intake levels. During experiments, consumption measurements occurred at 4 and 24 hours post-treatment, with corresponding 24-hour body weight changes documented. To assess tirzepatide’s impact on relapse-like behavior, we utilized the ADE model (25, 40). This protocol involved an 8-week baseline alcohol consumption period followed by 10-day alcohol deprivation within the intermittent access paradigm. Rats then received single tirzepatide or vehicle administration before alcohol reintroduction, with relapse-like drinking quantified as percentage change from baseline intake. A separate study examined repeated tirzepatide or vehicle administration effects spanning six alcohol drinking days across two weeks following the baseline period. This design allowed assessment of treatment efficacy over extended an timeframe. Upon completing the repeated administration study, rats were euthanized 24 hours after final treatment following a full day of alcohol access. Brains were rapidly removed, flash-frozen, and stored at -80°C for subsequent analysis. We additionally dissected and weighed several metabolically relevant tissues: gastrocnemius muscle, iBAT, sWAT, gWAT, rpWAT, and liver. The liver’s middle lobe was flash-frozen and stored at -80°C. Trunk blood was also collected using serum tubes (Z-gel tubes with clotting activator, Sarstedt, Germany) and stored at -80°C for later analysis.

### Drinking in the dark

A standardized DID protocol in adult male and female mice were used to examine binge-like alcohol drinking (39). The experimental design utilized repeated four-day alcohol access cycles, where days 1-3 involved two-hour sessions and day 4 extended to four hours as the primary test day. Three days without alcohol separated each cycle. Sessions started three hours after dark-phase onset, with food remaining available throughout while water was temporarily removed during alcohol access periods. The protocol timing required modification based on pharmacological considerations. Our initial intermittent access studies suggested that lower tirzepatide doses might not produce measurable effects within the standard four-hour assessment window, prompting this temporal adjustment to better capture potential treatment effects.

### Measurements of liver triglycerides and serum cytokines

Liver tissue samples from the repeated alcohol drinking experiment were processed for triglyceride analysis using standard lipid extraction techniques. Tissue was lysed in 2:1 chloroform:methanol and washed with 0.9 M NaCl to achieve phase separation, as previously described (85). The triglyceride-containing lower phase was collected and evaporated overnight. Dried triglyceride pellets were resuspended in isopropanol (Sigma Aldrich) and quantified using a commercial triglyceride kit (Randox Laboratories Ltd, Crumlin, UK) according to manufacturer’s instructions. Absorbance was measured at 500 nm with 546 nm correction using a Spectramax i3x multiplate reader (Molecular Devices, San Jose, CA, USA). Serum cytokine levels were

measured using a custom-ordered Bio-Plex Pro™ rat Cytokine Assay kit (10014905, Bio-Rad, Hercules, CA, USA) to quantify five cytokines, based on the literature (86, 87): IL-1β, IL-6, IL-10, TNFα, and MCP-1 using the Bio-Plex 200 system (Bio-Rad) according to manufacturer’s instructions.

### Electrophysiological recordings

Coronal brain slices (300 μm) were prepared 24 hours after tirzepatide or vehicle administration, as previously described (24, 88). Field potentials were evoked using a stimulation electrode positioned near (0.2-0.3 mm) the recording electrode in each brain region (NAc core/shell, mPFC, DLS, DMS, LS). Population spike (PS) amplitudes were evoked using seven-step increasing stimulation protocols. Paired-pulse stimulation (50 ms interpulse interval, 0.1 Hz) was used to calculate paired-pulse ratio (PPR, PS2/PS1), to estimate changes in the probability of transmitter release. Data were acquired using Clampfit 10.2 software (Molecular Devices, Foster City, CA, USA).

### Proteomics - Global relative quantification

The LS was microdissected using a brain-slicing matrix on dry ice, weighed, and stored at -80°C until analysis as previously described (24). Protein extraction employed lysis buffer (2% sodium dodecyl sulfate, 100 mM triethylammonium bicarbonate) with a Covaris ML230 ultrasonicator. Protein concentrations were determined using the Pierce BCA Protein Assay Kit (Thermo Scientific, Gothenburg, Sweden). Sample processing followed a modified SP3 method. Samples and references (40 μg) underwent reduction (100 mM DTT), alkylation (20 mM iodoacetamide), and precipitation on Sera-Mag™ SpeedBeads (Cytiva, Uppsala, Sweden) using ethanol. After washing and drying, beads were resuspended in 100 mM TEAB for protein digestion with Trypsin/Lys-C mix (1:25) for two hours, followed by trypsin (1:50) overnight. Following bead removal, peptide concentrations were determined using Pierce™ Quantitative Fluorometric Peptide Assay (Thermo Scientific). Peptide samples (20 μg) were labeled using TMT pro 18-plex isobaric mass tagging reagents (Thermo Fisher Scientific), pooled into one TMT-set, and purified using HiPPR Detergent Removal Resin and Pierce™ Peptide Desalting Spin Columns. The TMT-set underwent basic reversed-phase chromatography (bRP-LC, pH10) fractionation into 36 fractions over 70 minutes. Mass spectrometry analysis employed an Orbitrap Eclipse Tribrid mass spectrometer with FAIMS Pro ion mobility system interfaced with an nLC 1200 liquid chromatography system. Peptides were separated on a C18 35 cm column over 90 minutes, with data acquired using SPS MS3 methodology. Raw files were processed using Proteome Discoverer (Ver 3.0, Thermo Scientific) against UniProt Swiss-Prot Rattus norvegicus database using Sequest search engine. Only unique peptides were used for relative quantification, with proteins required to pass a 5% false discovery rate threshold. Proteins showing significant expression changes were cross-referenced with existing literature and the UniProtKB database (50) to identify those linked to histone and chromatin processes (complete reference list in **Table S1**).

### Statistics

Statistical analyses were performed using GraphPad Prism (version 10.4.1, GraphPad Software Inc., Boston, MA, USA). The statistical approach was as described previously (24). In brief, normal distribution was assessed using the Shapiro-Wilk test. All subsequent tests were two-tailed with significance threshold at p<0.05. For comparisons between two groups in behavioral, intake, or electrophysiology experiments, paired or unpaired Student’s t-tests were applied as appropriate. Comparisons among three or more groups employed one-way ANOVA with Bonferroni post-hoc tests. For experiments with repeated measures (microdialysis, repeated alcohol drinking, and electrophysiology), repeated-measures two-way ANOVA with Bonferroni post-hoc tests were utilized. Welch’s t-test was used on log2-transformed data to identify DEPs. Proteins with a p- value<0.05 and fold-change ≥10% (log2 fold-change ≤ -0.137 or ≥0.137) were considered as differentially expressed. All data are presented as mean ± standard error of the mean (SEM) with individual values shown when appropriate.

## ACKNOWLEDGEMENTS

The authors gratefully acknowledge the technical assistance and expertise of Ebba Frövenholt, Erika Lucente, Manisha Bergum Samad and Anna-Lena Leverin. Proteomic analysis was performed at the Proteomics Core Facility, Sahlgrenska academy, Gothenburg University, with financial support from SciLifeLab and BioMS. Animations were created using BioRender.

## Data availability

All data sets generated or analyzed during this study are available in the Source Data file. The MS proteomics data have been deposited to the ProteomeXchange Consortium (http://proteomecentral.proteomexchange.org) via the PRIDE partner repository with the data set identifier PXD063324. Any additional information required to reanalyze the data reported in this paper is available upon request from the corresponding author.

