## Supplementary Material for "Tirzepatide attenuates dopamine reward signaling and suppresses alcohol drinking and relapse-like behaviors in rodents"

**This PDF file includes:**

Figures S1 to S10
Tables S1

**Other supporting materials for this manuscript include the following:**

Dataset - Source Data File as excelfile
Table S1 - as excelfile

Fig. S1

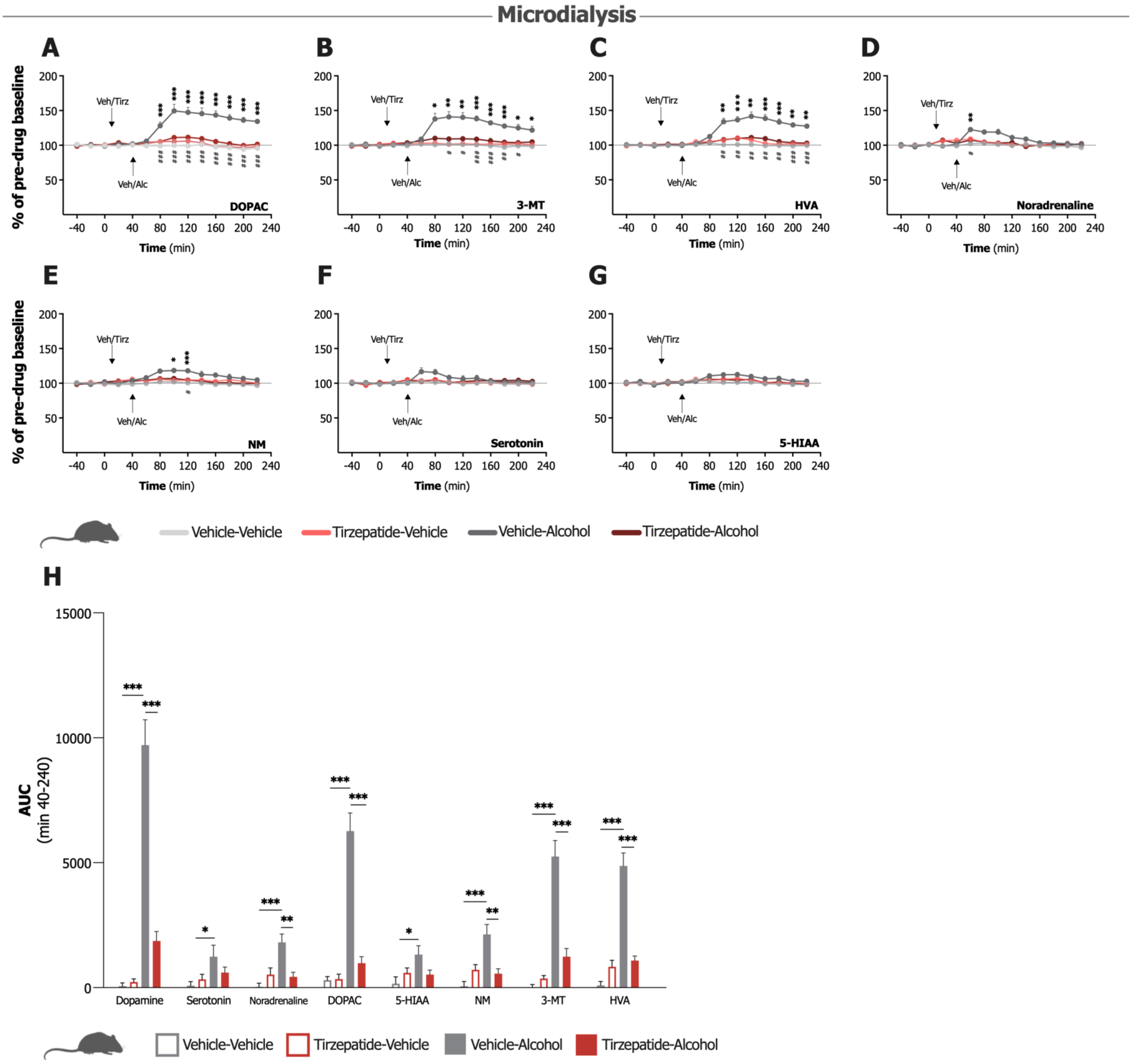

**Fig. S1. Accumbal monoamine levels in male mice following tirzepatide and systemic alcohol administration.**

**A-E.** Compared to vehicle (Veh), alcohol (Alc; 1.75 g/kg) administration evokes significant increases in the dopamine metabolites **(A)** 3,4-dihydroxyphenylacetic acid (DOPAC), **(B)** 3-methoxytyramine (3-MT), and **(C)** homovanillic acid (HVA) and **(D)** noradrenaline and **(E)** normetanephrine (NM) in the nucleus accumbens shell, effects that tirzepatide (Tirz; 0.144 mg/kg) treatment significantly attenuates (n=8/group, two-way repeated measures ANOVA). **F-G.** No significant effects occur in the levels of **(F)** serotonin and **(G)** 5-hydroxyindoleacetic acid (5-HIAA). **H.** Area under the curve (AUC) analysis confirms significant effects on dopamine, its metabolites (DOPAC, 3-MT, HVA), noradrenaline, and NM, as well as effects in alcohol-treated mice for serotonin and 5-HIAA, with more pronounced effects on monoamines related to dopamine (n=8/group, one-way ANOVA). Data show mean ± SEM. *P<0.05, **P<0.01, ***P<0.001, $^{\#}$P<0.05, $^{\#\#}$P<0.01, $^{\#\#\#}$P<0.001.

Fig. S2

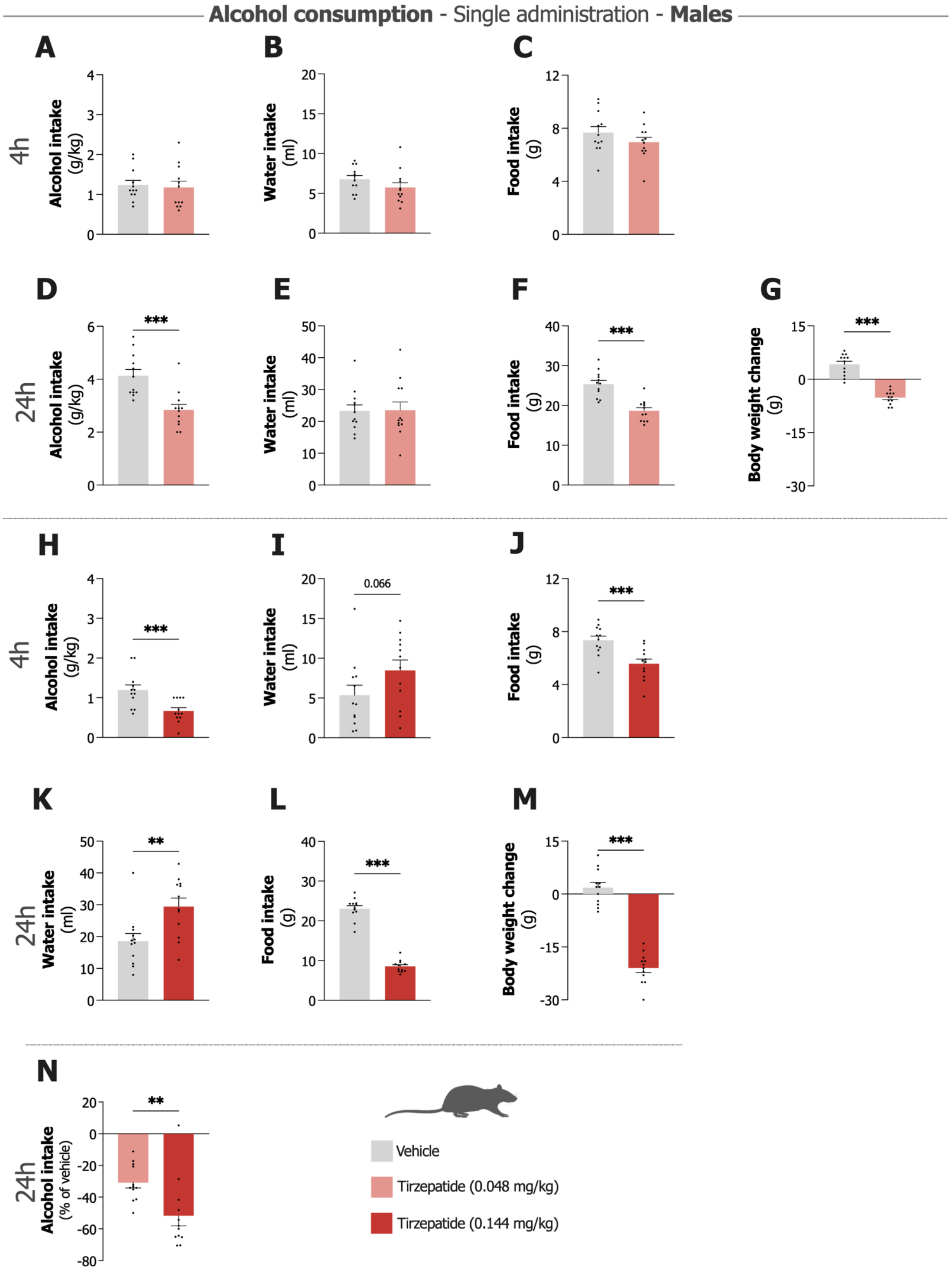

**Fig. S2. Effects of single administration of tirzepatide on intake of alcohol, water, and food, as well as body weight in male rats.**

**A-C.** Consumption at four hours following treatment with lower dose tirzepatide (0.048 mg/kg) shows no differences in **(A)** alcohol, **(B)** water, or **(C)** food intake compared to vehicle. **D-G.** At 24 hours the lower dose tirzepatide produces significant reductions in **(D)** alcohol and **(F)** food intake, along with decreases in **(G)** body weight compared to vehicle, while **(E)** water intake remains unaffected. **H-J.** Intake at four hours following treatment with the higher dose tirzepatide (0.144 mg/kg) shows reductions in **(H)** alcohol and **(J)** food intake, with a trend toward increased **(I)** water intake compared to vehicle. **K-M.** At 24 hours the higher dose increases **(K)** water intake and decreases **(L)** food intake and **(M)** body weight. **N.** Comparison of percent change in 24-hour alcohol intake against vehicle demonstrates the higher dose producing a more pronounced effect than the lower dose, indicating a dose-response effect. N=12/group, paired t-test. Data show individual data points with mean ± SEM. **P<0.01, ***P<0.001.

**Fig. S3**

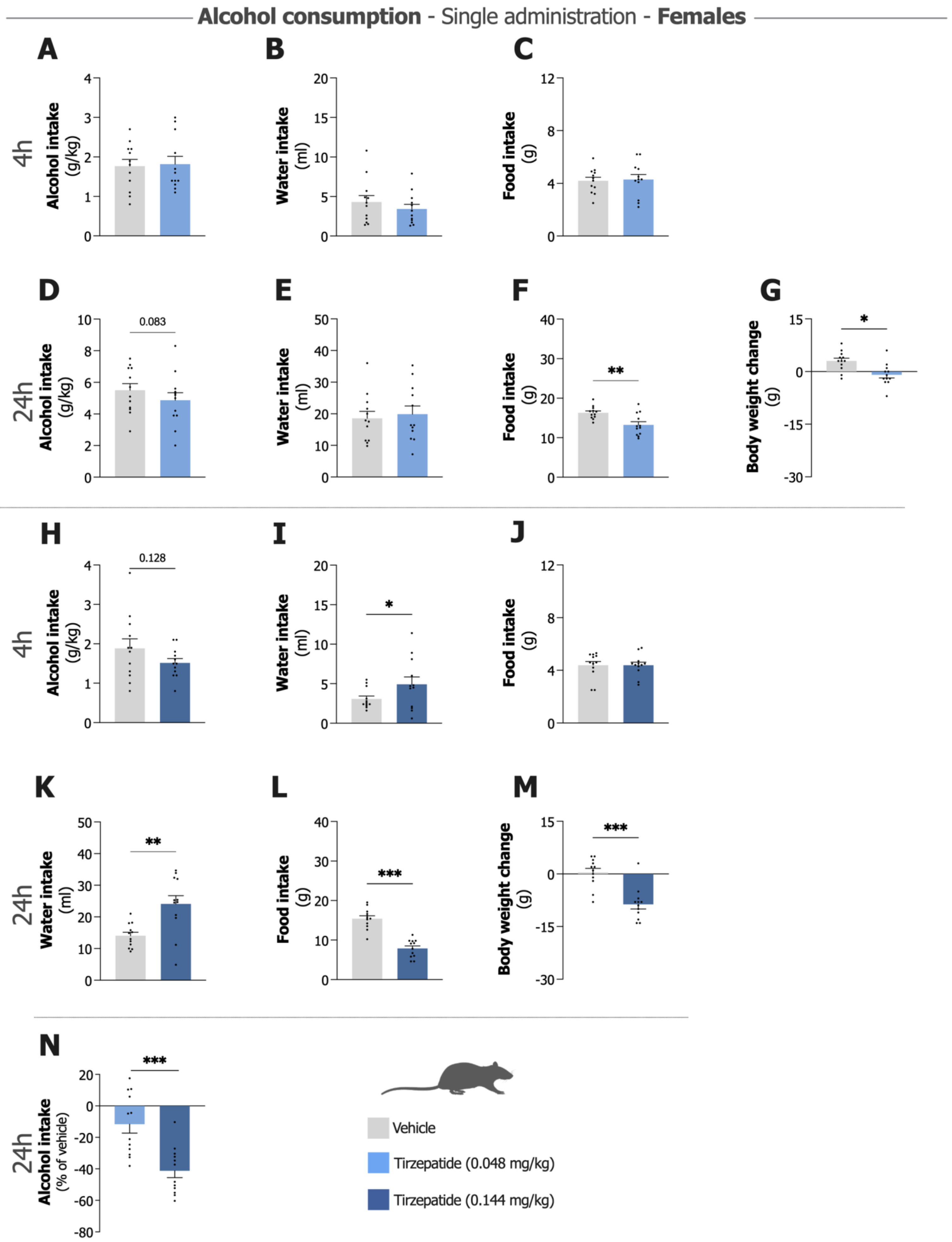

**Fig. S3. Effects of single administration of tirzepatide on intake of alcohol, water, and food, as well as body weight in female rats.**

**A-C.** Consumption at four hours following treatment with lower dose tirzepatide (0.048 mg/kg) shows no differences in **(A)** alcohol, **(B)** water, or **(C)** food intake compared to vehicle. **D-G.** At 24 hours the lower dose tirzepatide tends to reduce **(D)** alcohol intake, significantly decreases **(F)** food intake and **(G)** body weight compared to vehicle, while **(E)** water intake remains unaffected. **H-J.** Intake at four hours following treatment with the higher dose tirzepatide (0.144 mg/kg) demonstrates a trend to reduce **(H)** alcohol intake, shows an increase in **(I)** water intake, and produces no effect on **(J)** food intake compared to vehicle. **K-M.** At 24 hours the higher dose increases **(K)** water intake and decreases **(L)** food intake and **(M)** body weight compared to vehicle. **N.** Comparison of percent change in 24-hour alcohol intake against vehicle demonstrates the higher dose producing a more pronounced effect than the lower dose, indicating a dose-response effect. N=12/group, paired t-test. Data show individual data points with mean ± SEM. *P<0.05, **P<0.01, ***P<0.001.

**Fig. S4**

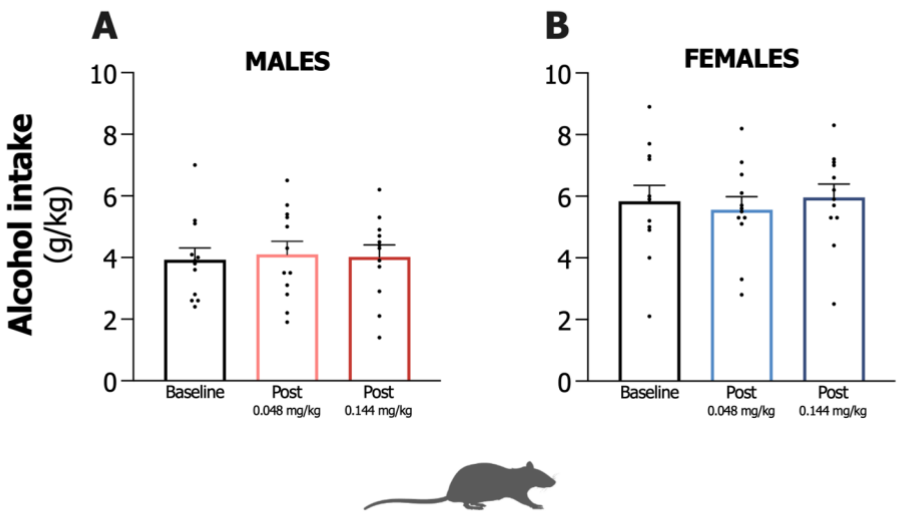

**Fig. S4. Single administration of tirzepatide does not produce persistent effects on alcohol intake in male and female rats.**

**A-B.** Alcohol consumption (g/kg) in **(A)** male and **(B)** female rats at baseline and 48 hours post-injection of tirzepatide at two doses (0.048 mg/kg and 0.144 mg/kg). Baseline alcohol intake and 48-hour post-injection levels show no differences at either dose, indicating a return to baseline drinking behavior, n=10 per group. Data show individual data points with mean ± SEM.

**Fig. S5**

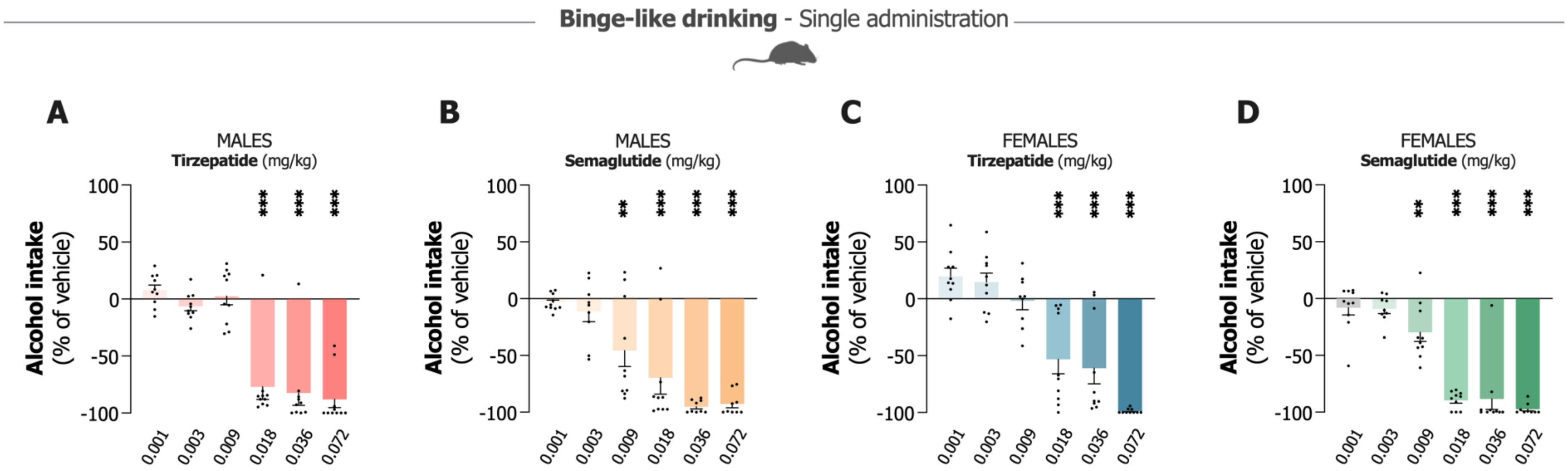

**Fig. S5. Dose-dependent effects of tirzepatide and semaglutide on binge-like alcohol drinking in male and female mice.**

**A-B.** Dose effects in male mice showing percent change in alcohol intake relative to vehicle following single administration of **(A)** tirzepatide or **(B)** semaglutide (0.001-0.072 mg/kg, intraperitoneally (IP)). Both drugs produce significant dose-dependent reductions in alcohol consumption, n=9-10/group, one-way ANOVA. **C-D.** Parallel experiments in female mice demonstrate dose-dependent decreases in alcohol intake following **(C)** tirzepatide or **(D)** semaglutide administration IP. Both compounds significantly reduce binge-like alcohol consumption, n=9-10/group, one-way ANOVA. Data show individual data points with mean ± SEM. Statistical significance compared to vehicle: **P<0.01, ***P<0.001.

**Fig. S6**

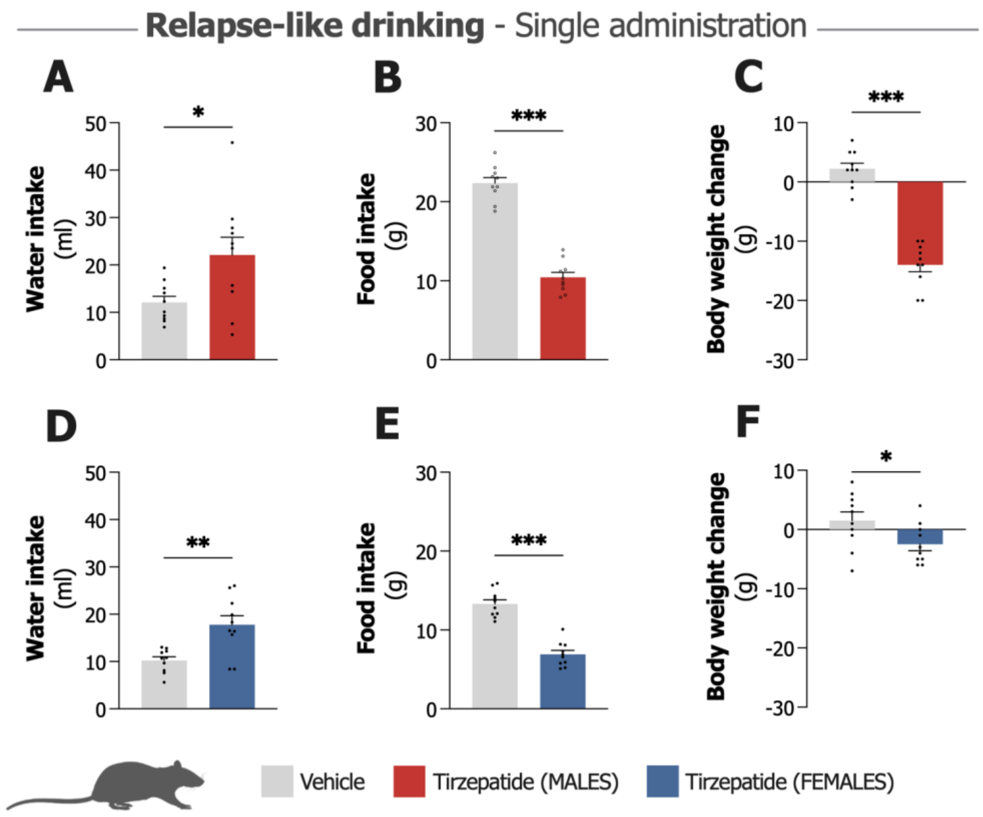

**Fig. S6. Effects of single dose tirzepatide on water intake, food intake, and body weight in the relapse-like drinking paradigm in male and female rats.**

**A-C.** Intake in male rats at 24 hours following single tirzepatide treatment shows significant increases in **(A)** water intake, decreases in **(B)** food intake, and **(C)** body weight compared to vehicle. **D-F.** Examination of the same parameters in female rats reveals increases in **(D)** water intake and decreases in **(E)** food intake and **(F)** body weight compared to vehicle. N=10/group, paired t-test. Data show individual data points with mean ± SEM. *P<0.05, **P<0.01, ***P<0.001.

**Fig. S7**

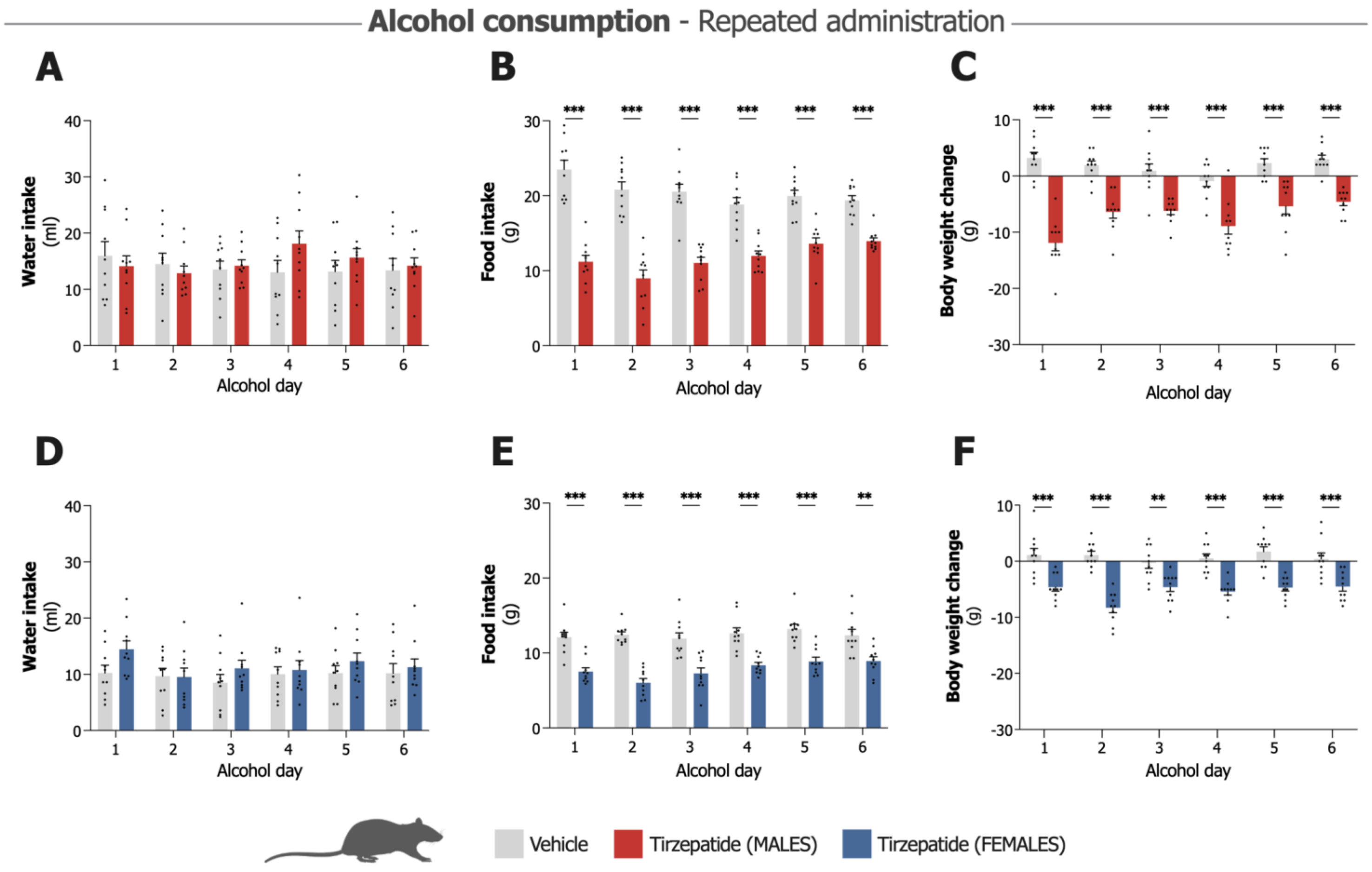

**Fig. S7. Effects of repeated tirzepatide administration on water intake, food intake, and body weight in male and female rats.**

**A-C. (A)** Water intake remains unaffected in male rats across all six alcohol drinking days compared to vehicle, while **(B)** food intake and **(C)** body weight are reduced by tirzepatide treatment. **D-F.** Examination of the same parameters in female rats reveals no significant changes in **(D)** water intake following tirzepatide treatment. **(E)** Food intake and **(F)** body weight show significant reductions throughout the paradigm, compared to vehicle administration. N=10/group, paired t-test. Data show individual data points with mean ± SEM. **P<0.01, ***P<0.001.

**Fig. S8**

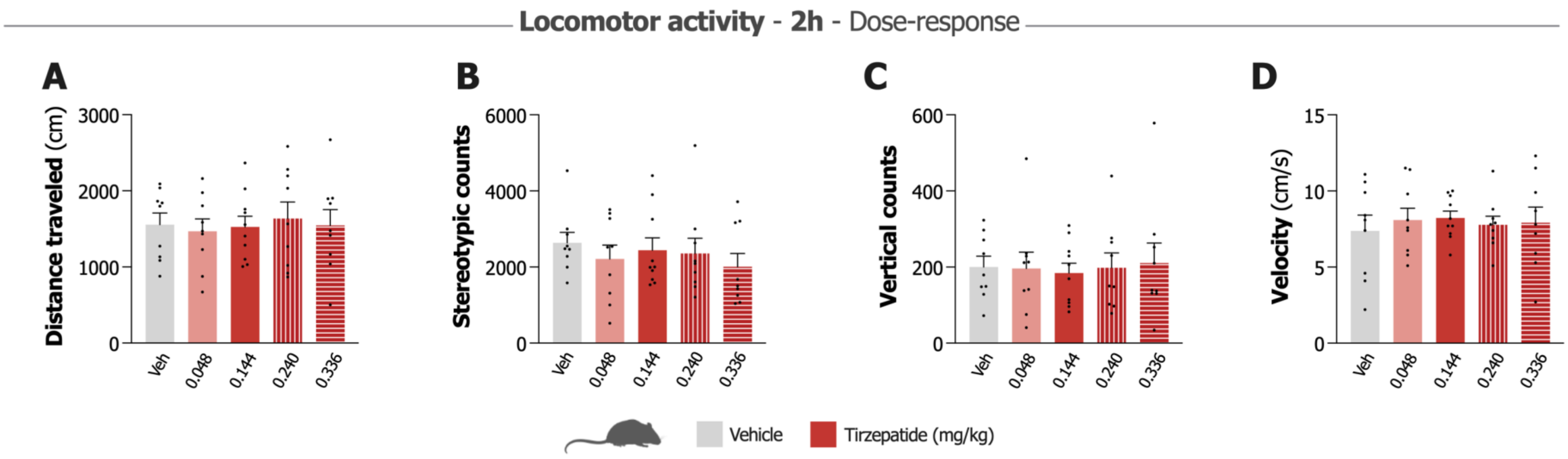

**Fig. S8. Dose-response effects of tirzepatide on locomotor activity in male mice.**

**A-D.** Dose-response effects of tirzepatide on locomotor activity over two hours show no significant differences in distance traveled **(A)**, stereotypic counts **(B)**, vertical counts **(C)**, or velocity **(D)** across all doses compared to vehicle (Veh; n=9-10/group, one-way ANOVA). Data show individual data points with mean ± SEM.

**Fig. S9**

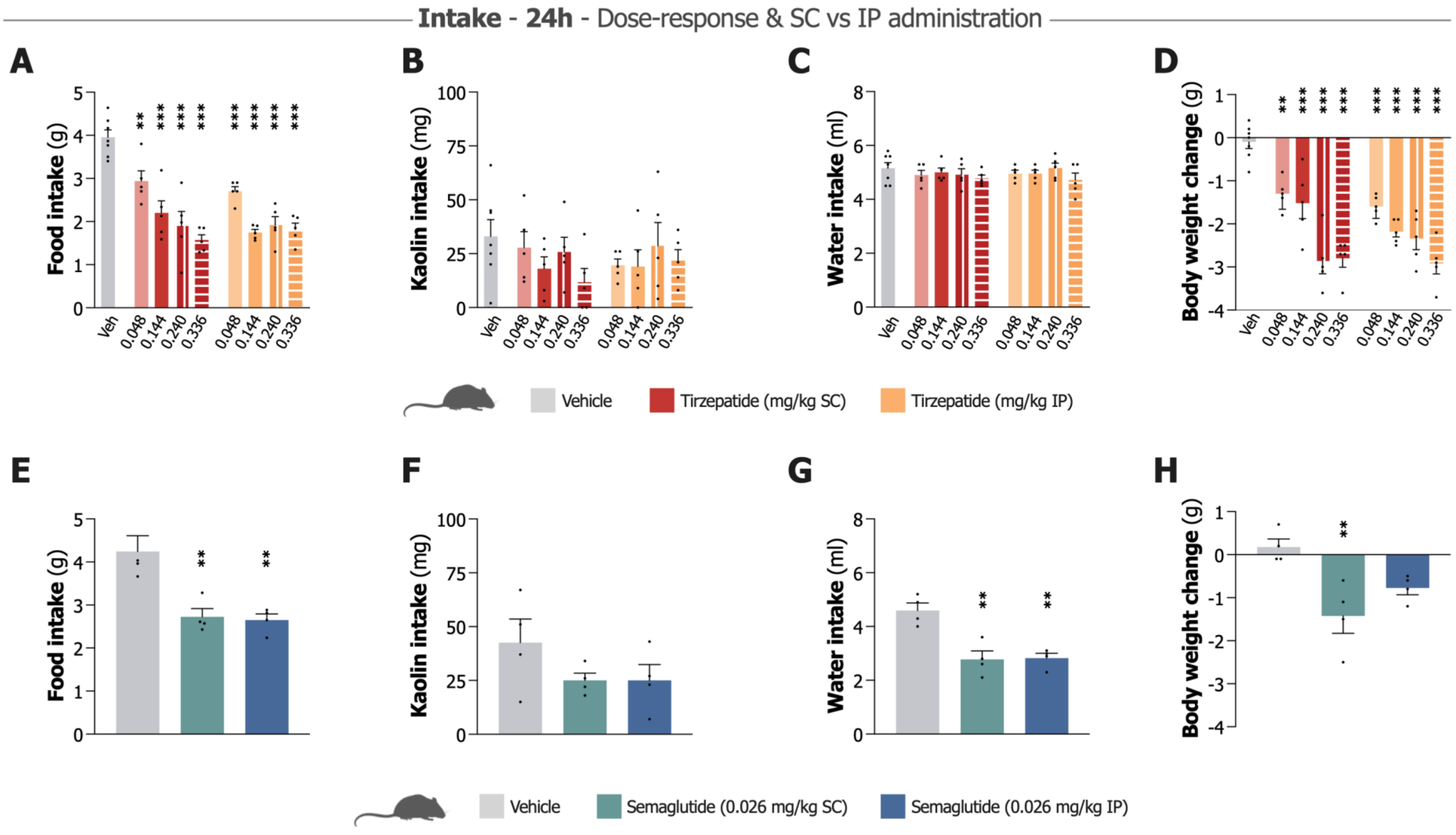

**Fig. S9. Effects of tirzepatide and semaglutide on intake of food, water and kaolin, and body weight through different routes of administration in male mice.**

Baseline consumption was recorded for three consecutive days prior to these pilot experiments. On the test day, both drugs were administered one hour before the onset of the dark cycle, with all subsequent measurements conducted 24 hours later. **A-D.** Dose-response effects of tirzepatide administered via intraperitoneal (IP) or subcutaneous (SC) injection on 24-hour intake measurements. Tirzepatide significantly reduces food intake **(A)** at all doses tested via both routes compared to vehicle. No significant effects occur on water **(B)** or kaolin intake **(C)**. Body weight change **(D)** shows significant reductions at all doses via both routes compared to vehicle (n=5/group, one-way ANOVA). **E-H.** Effects of semaglutide (0.026 mg/kg) administered via SC or IP injection on 24-hour intake measurements. Both routes of administration significantly reduce food **(E)** and water intake **(F)** compared to vehicle. No significant effects occur on kaolin intake **(G)**. Body weight change **(H)** shows significant reductions with both routes of administration compared to vehicle (n=4/group, one-way ANOVA). Data show individual data points with mean ± SEM. ***P<0.001, **P<0.01.

**Fig. S10**

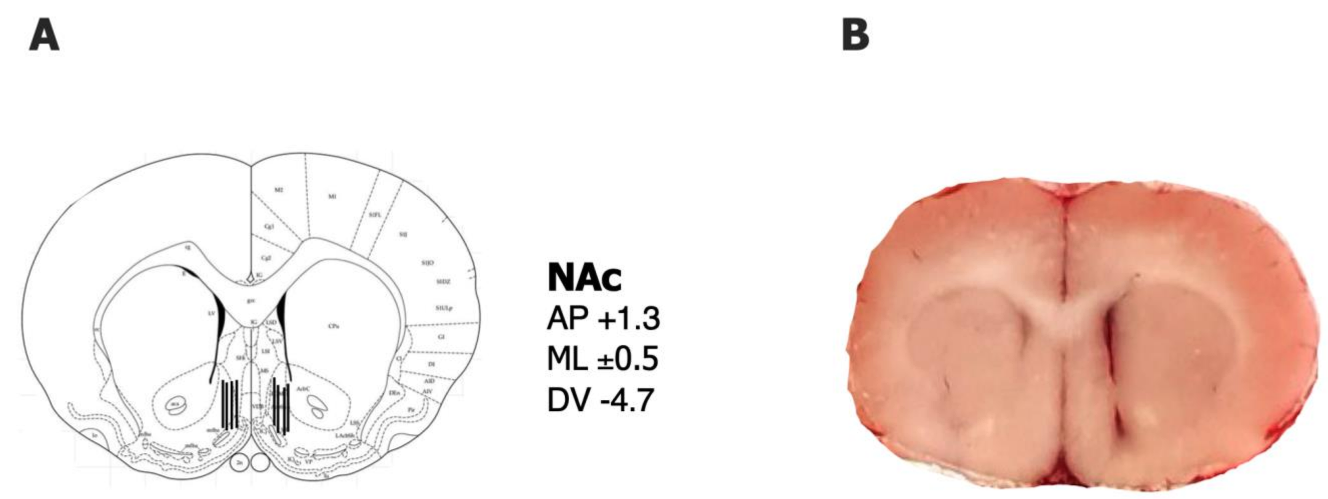

**Fig. S10. Probe placements.**

**A.** Schematic representation of microdialysis probe placements within the nucleus accumbens (NAc) shell, including stereotaxic coordinates. **B.** A representative placement of a microdialysis probe targeting NAc shell in male mice.

### Table S1

| Accession | Gene symbol | Protein description | p-value | -log10 (p-value) | log2 (fold change) | Biological function or pathway | Reference (UniProtKB) | Additional references |
|---|---|---|---|---|---|---|---|---|
| Q4V8E5 | MFSD3 | Major facilitator superfamily domain-containing protein 3 | 5,75E-03 | 2,24 | 0,336 | Transporter activity | https://www.ebi.ac.uk/QuickGO/annotations?geneProductId=Q4V8E5 | |
| P43278 | H1-0 | Histone H1.0 | 1,38E-02 | 1,86 | 0,333 | Histone linker | https://www.ebi.ac.uk/QuickGO/annotations?geneProductId=P43278 | |
| Q496Z9 | TRMT1L | TRMT1-like protein | 6,92E-03 | 2,16 | 0,332 | Adult behavior, role in motor coordination | https://www.ebi.ac.uk/QuickGO/annotations?geneProductId=Q496Z9 | |
| P15865 | H1-4 | Histone H1.4 | 2,40E-02 | 1,62 | 0,326 | Histone linker | https://www.ebi.ac.uk/QuickGO/annotations?geneProductId=P15865 | |
| O08701 | ARG2 | Arginase-2, mitochondrial | 3,98E-02 | 1,40 | 0,304 | Glucose homeostasis, insulin sensitivity | https://www.ebi.ac.uk/QuickGO/annotations?geneProductId=O08701 | |
| Q9WVK3 | PECR | Peroxisomal trans-2-enoyl-CoA reductase | 5,62E-03 | 2,25 | 0,266 | Fatty acid biosynthesis, Neuroinflammation, Alcohol related effects | https://www.ebi.ac.uk/QuickGO/annotations?geneProductId=Q9WVK3 | |
| Q9R1S9 | MDK | Midkine | 2,09E-02 | 1,68 | 0,248 | Brain development, regulate ethanol consumption, behavior, short-term memory | https://www.ebi.ac.uk/QuickGO/annotations?geneProductId=Q9R1S9 | |
| P13437 | ACAA2 | 3-ketoacyl-CoA thiolase, mitochondrial | 2,57E-02 | 1,59 | 0,244 | Fatty acid biosynthesis, associated with alcohol use disorder | https://www.ebi.ac.uk/QuickGO/annotations?geneProductId=P13437 | |
| Q4KM93 | TMEM177 | Transmembrane protein 177 | 2,88E-02 | 1,54 | 0,239 | Mitochondrial respiratory chain complex I assembly | https://www.ebi.ac.uk/QuickGO/annotations?geneProductId=Q4KM93 | |
| P60889 | CBX7 | Chromobox protein homolog 7 | 7,41E-03 | 2,13 | 0,238 | Chromatin binding, methylated histone, promotes histone H3 trimethylation (H3K9me3) | https://www.ebi.ac.uk/QuickGO/annotations?geneProductId=P60889 | doi: 10.1038/s41467-022-34269-7 , doi: 10.1038/s41418-018-0064-0 |
| Q05096 | MYO1B | Unconventional myosin-Ib | 3,39E-02 | 1,47 | 0,232 | Nervous system development | https://www.ebi.ac.uk/QuickGO/annotations?geneProductId=Q05096 | |
| Q499T2 | IFI30 | Gamma-interferon-inducible lysosomal thiol reductase | 3,80E-02 | 1,42 | 0,231 | Immune and protein stabilization | https://www.ebi.ac.uk/QuickGO/annotations?geneProductId=Q499T2 | |
| Q5FVM1 | TMEM231 | Transmembrane protein 231 | 3,47E-03 | 2,46 | 0,228 | Regulation of protein localization | https://www.ebi.ac.uk/QuickGO/annotations?geneProductId=Q5FVM1 | |
| B2GV91 | LYRM2 | LYR motif-containing protein 2 | 3,63E-02 | 1,44 | 0,224 | Mitochondrial respiratory chain complex I assembly | https://www.ebi.ac.uk/QuickGO/annotations?geneProductId=B2GV91 | |
| A1L1L7 | ZNF426 | Zinc finger protein 426 | 7,59E-04 | 3,12 | 0,222 | DNA processes and RNA | https://www.ebi.ac.uk/QuickGO/annotations?geneProductId=A1L1L7 | |
| Q5RKI3 | POLL | DNA polymerase lambda | 1,45E-02 | 1,84 | 0,222 | DNA processes | https://www.ebi.ac.uk/QuickGO/annotations?geneProductId=Q5RKI3 | |
| P18437 | HMGN2 | Non-histone chromosomal protein HMG-17 | 4,17E-03 | 2,38 | 0,221 | Chromatin organization | https://www.ebi.ac.uk/QuickGO/annotations?geneProductId=P18437 | doi: 10.1186/s13072-019-0320-7 , doi: 10.1074/jbc.M114.555425 |
| P31643 | SLC6A6 | Sodium- and chloride-dependent taurine transporter | 6,17E-03 | 2,21 | 0,219 | GABA-ergic synapse, GABA import, neurotransmitter, taurine | https://www.ebi.ac.uk/QuickGO/annotations?geneProductId=P31643 | |
| P31503 | POU2F1 | POU domain, class 2, transcription factor 1 | 4,90E-02 | 1,31 | 0,194 | Chromatin binding, activates the promoters of genes such as those for histone H2B | https://www.ebi.ac.uk/QuickGO/annotations?geneProductId=P31503 | doi: 10.1016/S0021-9258(19)74567-0 , doi: 10.1093/nar/gky635 |
| A6N6J5 | WDR35 | WD repeat-containing protein 35 | 4,17E-02 | 1,38 | 0,193 | Cellular response to glucose stimulus | https://www.ebi.ac.uk/QuickGO/annotations?geneProductId=A6N6J5 | |
| Q9Z1B2 | GSTM5 | Glutathione S-transferase Mu 5 | 6,46E-05 | 4,19 | 0,178 | Glutathione metabolic process | https://www.ebi.ac.uk/QuickGO/annotations?geneProductId=Q9Z1B2 | |
| P36860 | RALB | Ras-related protein Ral-B | 1,70E-02 | 1,77 | 0,178 | Oncogen, apoptotic processis, response to starvation | https://www.ebi.ac.uk/QuickGO/annotations?geneProductId=P36860 | |
| Q4V7F5 | PIH1D1 | PIH1 domain-containing protein 1 | 9,12E-03 | 2,04 | 0,177 | Regulation of glucose signaling, mTORC, histone binding, chromatin remodelling | https://www.ebi.ac.uk/QuickGO/annotations?geneProductId=Q4V7F5 | doi: 10.1016/j.celrep.2014.03.013 , doi: 10.1093/jmcb/mjs003 |
| Q4KLZ6 | TKFC | Triokinase/FMN cyclase | 3,47E-02 | 1,46 | 0,174 | Fructose catabolism/metabolism | https://www.ebi.ac.uk/QuickGO/annotations?geneProductId=Q4KLZ6 | |
| Q641Y9 | DALRD3 | DALR anticodon-binding domain-containing protein 3 | 2,69E-02 | 1,57 | 0,162 | tRNA processes | https://www.ebi.ac.uk/QuickGO/annotations?geneProductId=Q641Y9 | |
| Q9JIM4 | DUSP12 | Dual specificity protein phosphatase 12 | 5,37E-03 | 2,27 | 0,161 | Chromatin binding and remodelling, histone H2AXS140 and H2AXS142 phosphatase activity | https://www.ebi.ac.uk/QuickGO/annotations?geneProductId=Q9JIM4 | doi: 10.1016/j.jprot.2019.02.008 |
| Q80W57 | ABCG2 | Broad substrate specificity ATP-binding cassette transporter CG2 | 9,55E-03 | 2,02 | 0,156 | Response to alcohol, response to insulin lipid homeostasis | https://www.ebi.ac.uk/QuickGO/annotations?geneProductId=Q80W57 | |
| Q4QQT6 | BRIX1 | Ribosome biogenesis protein BRX1 homolog | 2,51E-02 | 1,60 | 0,155 | rRNA processing | https://www.ebi.ac.uk/QuickGO/annotations?geneProductId=Q4QQT6 | |
| Q5RKH7 | SLC35F6 | Solute carrier family 35 member F6 | 4,47E-02 | 1,35 | 0,153 | Transmembrane transport | https://www.ebi.ac.uk/QuickGO/annotations?geneProductId=P31643 | |
| P84245 | H3-3B | Histone H3.3 | 4,79E-03 | 2,32 | 0,149 | Histone | https://www.ebi.ac.uk/QuickGO/annotations?geneProductId=P84245 | |
| P02262 | H2A-1A | Histone H2A type 1 | 6,92E-03 | 2,16 | 0,149 | Histone | https://www.ebi.ac.uk/QuickGO/annotations?geneProductId=P02262 | |
| Q642A6 | VWA1 | von Willebrand factor A domain-containing protein 1 | 4,47E-02 | 1,35 | 0,141 | Behavioral response to pain, regulation of Insulin-like Growth Factor (IGF) transport | https://www.ebi.ac.uk/QuickGO/annotations?geneProductId=Q642A6 | |
| Q8K4H4 | APTX | Aprataxin | 4,37E-02 | 1,36 | 0,139 | DNA repair, chromatin | https://www.ebi.ac.uk/QuickGO/annotations?geneProductId=Q8K4H4 | doi: 10.1093/nar/gky507 , doi: 10.1093/hmg/ddh122 |
| P57790 | KEAP1 | Kelch-like ECH-associated protein 1 | 4,68E-02 | 1,33 | 0,138 | Regulator of Nrf2, oxidative stress | https://www.ebi.ac.uk/QuickGO/annotations?geneProductId=P57790 | |
| Q6P798 | RCBTB2 | RCC1 and BTB domain-containing protein 2 | 3,63E-02 | 1,44 | 0,137 | Liver and GI development | https://www.ebi.ac.uk/QuickGO/annotations?geneProductId=Q6P798 | |
| P97710 | SIRPA | Tyrosine-protein phosphatase non-receptor type substrate 1 | 3,31E-02 | 1,48 | -0,586 | Response to interleukin, synaptic loss, neurodegeneration, neuroinflammation | https://www.ebi.ac.uk/QuickGO/annotations?geneProductId=P97710 | |
| Q5BK56 | GSTM4 | Glutathione S-transferase Mu 4 | 1,32E-03 | 2,88 | -0,519 | Glutathione metabolic process, lipid metabolic processes | https://www.ebi.ac.uk/QuickGO/annotations?geneProductId=Q5BK56 | |
| Q9QZQ5 | CCN3 | CCN family member 3 | 3,31E-02 | 1,48 | -0,352 | Regulation of insulin secretion, regulation of Notch signaling pathway | https://www.ebi.ac.uk/QuickGO/annotations?geneProductId=Q9QZQ5 | |
| O35815 | ATXN3 | Ataxin-3 | 1,10E-02 | 1,96 | -0,209 | Exploration behavior, mTORC1 signaling (linked to alcohol histone deacetylase activity | https://www.ebi.ac.uk/QuickGO/annotations?geneProductId=O35815 | doi: 10.1093/nar/gkad212 , doi: 10.1523/JNEUROSCI.2053-06.2006 |
| Q5BJT4 | TXNDC15 | Thioredoxin domain-containing protein 15 | 3,09E-02 | 1,51 | -0,207 | Tyrosine biosynthetic process, by oxidation of phenylalanine | https://www.ebi.ac.uk/QuickGO/annotations?geneProductId=Q5BJT4 | |
| Q6RJR6 | RTN3 | Reticulon-3 | 2,82E-02 | 1,55 | -0,204 | Brain development, glutamatergic synapse, obesity triglycerides | https://www.ebi.ac.uk/QuickGO/annotations?geneProductId=Q6RJR6 | |
| Q6AXT7 | RBM42 | RNA-binding protein 42 | 1,32E-02 | 1,88 | -0,201 | mRNA binding activity/mRNA splicing | https://www.ebi.ac.uk/QuickGO/annotations?geneProductId=Q6AXT7 | |
| Q569B5 | LRRC14 | Leucine-rich repeat-containing protein 14 | 1,51E-02 | 1,82 | -0,198 | Toll-like receptor | https://www.ebi.ac.uk/QuickGO/annotations?geneProductId=Q569B5 | |
| Q64548 | RTN1 | Reticulon-1 | 3,80E-02 | 1,42 | -0,198 | Brain development, alcohol-regulated gene | https://www.ebi.ac.uk/QuickGO/annotations?geneProductId=Q64548 | |
| D3ZFB6 | PRRT2 | Proline-rich transmembrane protein 2 | 3,16E-02 | 1,50 | -0,194 | Glutamatergic synapse, post- and presynapse, movement disorders | https://www.ebi.ac.uk/QuickGO/annotations?geneProductId=D3ZFB6 | |
| A2VCW9 | AASS | Alpha-aminoadipic semialdehyde synthase, mitochondrial | 4,17E-02 | 1,38 | -0,190 | Saccharopine dehydrogenase (NAD+, L-glutamate-forming) activity,glutamate learning | https://www.ebi.ac.uk/QuickGO/annotations?geneProductId=A2VCW9 | |
| B1WBU8 | PLEKHD1 | Pleckstrin homology domain-containing family D member 1 | 4,07E-02 | 1,39 | -0,175 | | https://www.uniprot.org/uniprotkb/B1WBU8/entry | |
| M0RDU0 | F8A1 | 40-kDa huntingtin-associated protein | 2,63E-02 | 1,58 | -0,169 | Regulation of proteasomal protein catabolic process , vesicle cytoskeletal trafficking | https://www.ebi.ac.uk/QuickGO/annotations?geneProductId=M0RDU0 | |
| O88339 | EPN1 | Epsin-1 | 4,07E-02 | 1,39 | -0,151 | Post- and presynapse, Notch signaling pathway | https://www.ebi.ac.uk/QuickGO/annotations?geneProductId=O88339 | |
| Q01460 | CTBS | Di-N-acetylchitobiase | 4,57E-02 | 1,34 | -0,139 | Oigosaccharide catabolic process | https://www.ebi.ac.uk/QuickGO/annotations?geneProductId=Q01460 | |
| P34926 | MAP1A | Microtubule-associated protein 1A | 5,25E-03 | 2,28 | -0,137 | Glutamatergic synapse, memory, associative learning, glutamate, synatic plasticity | https://www.ebi.ac.uk/QuickGO/annotations?geneProductId=P34926 | |

**Table S1. All differentially expressed proteins and their detailed gene ontology-annotations and reactome pathway.**

Includes all differentially expressed proteins (DEP)s and their detailed gene ontology (GO)-annotations and reactome pathway, p-value 0.05 and Fold change Log2 ±0.137
